# Gamma-Oscillation Plasticity Is Mediated by Parvalbumin Interneurons

**DOI:** 10.1101/2023.06.21.545901

**Authors:** Michael D. Hadler, Alexandra Tzilivaki, Dietmar Schmitz, Henrik Alle, Jörg R. P. Geiger

**Affiliations:** Charité-Universitätsmedizin Berlin, corporate member of Freie Universität Berlin, Humboldt-Universität zu Berlin, Berlin Institute of Health, Charitéplatz 1, 10117 Berlin, Germany; Institute of Neurophysiology, Charité-Universitätsmedizin Berlin, Berlin, Germany; Einstein Center for Neurosciences Berlin, Charitéplatz 1, 10117 Berlin, Germany; Neurocure Cluster of Excellence, Charitéplatz 1, 10117 Berlin, Germany; German Center for Neurodegenerative Diseases (DZNE), Berlin, Germany; Bernstein Center for Computational Neuroscience, Berlin, Germany; Max Delbrück Center for Molecular Medicine in the Helmholtz Association, Robert-Rössle-Straße 10, 13125 Berlin, Germany

## Abstract

Understanding the plasticity of neuronal networks is an emerging field of (patho-)physiological research, yet little is known about the underlying cellular mechanisms. Gamma-oscillations (30 – 80 Hz), a biomarker of cognitive performance, require and potentiate glutamatergic transmission onto parvalbumin-positive interneurons (PVIs), suggesting an interface for cell-to-network plasticity. In *ex vivo* local field potential recordings, we demonstrate long-term potentiation of hippocampal gamma-power. Gamma-potentiation obeys established rules of PVI plasticity, requiring calcium-permeable AMPA receptors (CP-AMPARs) and metabotropic glutamate receptors (mGluRs). A microcircuit model of CA3 gamma-oscillations predicts CP-AMPAR plasticity onto PVIs critically outperforms pyramidal cell plasticity in increasing gamma-power and completely accounts for gamma-potentiation. We re-affirm this ex vivo in three PVI-targeting animal models, demonstrating that gamma-potentiation requires PVI-specific metabotropic signaling via a Gq/PKC-pathway comprising mGluR5 and a Gi-sensitive, PKA-dependent pathway. Gamma-activity dependent, metabotropically mediated CP-AMPAR plasticity on PVIs may serve as a guiding principle in understanding network plasticity in health and disease.

## Introduction

Cortical networks are thought to implement task-specific computations by synchronizing the firing patterns of neurons to a set of defined rhythms^1, 2^. Neuronal oscillations are distinct regarding both the cortical state they accompany as well as the underlying synaptic interactions between the participating neurons^3^. Similar to the plasticity of synaptic weights, the spectral profiles of distinct oscillatory patterns adapt following learning^4–6^, or deteriorate in states of disease^7^, expressed as changes in their spectral amplitudes and/or frequency distributions. Crucially, such changes coincide with either beneficial or detrimental changes in the respective behavioral or cognitive performance. Experimental approaches utilizing either sensory stimuli or optogenetic strategies corroborate the causal link between oscillatory amplitude and performance on short time scales yet fail to explain how these changes are recalled after prolonged periods as required for successful learning^8–10^. This warrants a cellular storage mechanism of oscillatory response tuning innate to neuronal networks.

Hippocampal gamma-oscillations (30 – 80 Hz) contribute to the generation, storage and retrieval of memories and have been studied extensively *in vivo*^11, 12^, *ex vivo*^13^ and *in silico*^14^. In the CA3 subregion, an understanding has emerged that gamma-activity results from precisely-timed synaptic feedback loops between local pyramidal cells and interneurons^15, 16^. Particularly fast-spiking, parvalbumin-positive interneurons (PVIs) are equipped with specific synaptic properties that facilitate synchronization at gamma-frequencies, as they quickly transform converging glutamatergic inputs via GluA2-lacking, calcium-permeable AMPA receptors (CP-AMPARs) into divergent, powerful inhibition^17^. During periods of increased neuronal activity, this promotes the co-activation of postsynaptic cells at short time intervals^18, 19^ and benefits the induction of synaptic plasticity. In line with this, recent studies have highlighted a vital importance of PVI activation for memory formation and maintenance^20–23^. Yet it is unclear how this relates to their role in promoting synchrony and further complicated by the fact that PVIs themselves are subjected to various forms of anatomical^24^, molecular^25^ and synaptic plasticity^26^. It is therefore conceivable that PVI plasticity is sufficient to store long-term changes of gamma-activity, facilitating its reinstatement upon retrieval.

We previously demonstrated that the induction of gamma-oscillations *in vivo* and *ex vivo* evokes a long-lasting plasticity of sharp-wave ripple complexes, which is accompanied by long-term potentiation (LTP) of glutamatergic inputs onto both pyramidal cells and PVIs^27^. Gamma-induced plasticity is mediated via group I metabotropic glutamate receptors (mGluRs) and largely independent of NMDA receptor (NMDAR) activation, yet a broader profile of the involved signaling cascades remains to be determined. This prompts the question whether these acute plastic changes on a synaptic level affect later overall network activity and if so, which role is attributed to the specific plasticity obtained by PVIs. Here, in an *ex vivo* slice model of murine CA3, we demonstrate that evoked gamma-power is markedly increased hours after previous episodes of gamma-activity. CP-AMPARs are required not only for the generation of the gamma-rhythm, but also mediate the subsequent increase of power we term “gamma-potentiation”. In an *in silico* microcircuit model of CA3 gamma-oscillations, we predict that an increase of CP-AMPAR conductances at the pyramidal cell-to-PVI synapse completely accounts for gamma-potentiation. Using both pharmacological and genetic tools specific to known plasticity rules of PVIs, we confirm that gamma-potentiation can be explained entirely by the activation of metabotropic pathways in PVIs and uncover an additional requirement of both PKC and PKA activation in PVIs. The transfer of PV input plasticity to output oscillations provides a synaptic basis to gamma frequency-specific network plasticity.

## Results

### Gamma-potentiation in the mouse hippocampus

In acute transverse hippocampal slices of adolescent mice (p45 – p70) we performed either multiple single-site local field potential (LFP) or perforated multi-electrode array (pMEA) recordings in the CA3 pyramidal cell layer. After an initial recording of baseline activity in LFP recordings, in which no oscillatory activity was detected, network oscillations in the low-gamma frequency range (25 – 45 Hz, recorded at 32 – 34° C) were reliably induced by bath-application of kainate (KA1, 150 nM for 30 minutes at 1.5 ml/min). Following a one-hour washout period, during which oscillatory activity had completely subsided after 30 – 40 minutes, a second, identical period of network activity was induced (KA2, 150 nM for 30 minutes). During this second application period of KA, peak low-gamma power on average increased approximately two-fold (**Fig. 1A-C**, KA1: 5.66 [1.88 16.93] µV² vs. KA2: 11.84 [3.40 32.50] µV², n = 15 slices tested, Wilcoxon Signed Rank Test: p = 6.1 × 10^-5^). Whereas the absolute values of peak gamma-power and frequency in the first induction period were highly variable across individual slices and conditions tested (**Supplementary Fig. 1**), they did not correlate with the subsequent relative increase of peak power (Power KA2/KA1: 2.13 +/- 0.11, from here on referred to as gamma-potentiation). Across all sole LFP experiments reported in this study, the average potentiation was highly reproducible amongst control conditions across experimental groups, highlighting the stability of our *ex vivo* approach (**Supplementary Fig. 1**). We further confirmed that the increase of peak power was conserved even if the washout period was extended to three hours, suggesting a long-lasting change of oscillatory network excitability (**Fig. 1B-C**, Power KA2/KA1: 2.99 +/- 0.71, n = 10). Potentiation was concomitantly observed in down-stream CA1 (**Supplementary Fig. 2a,c,** Power KA2/KA1: 2.80 +/- 0.52, n = 15), which in intact slices is synchronized by CA3 in the low-gamma frequency range via the Schaffer collateral pathway^28^. However, in “CA1-Mini” slices, in which CA3 and the subiculum are disconnected from CA1^29^, application of KA evokes rhythmic mid-gamma activity (50 – 60 Hz, recorded at 32 – 34° C). Applying an adjusted application protocol (400 nM KA, 2 × 30 minutes with 60 minutes wash-out) again revealed a roughly two-fold increase of peak mid-gamma power (**Supplementary Fig. 2b,d,** Power KA2/KA1: 1.81 +/- 0.12).

**Figure 1.**
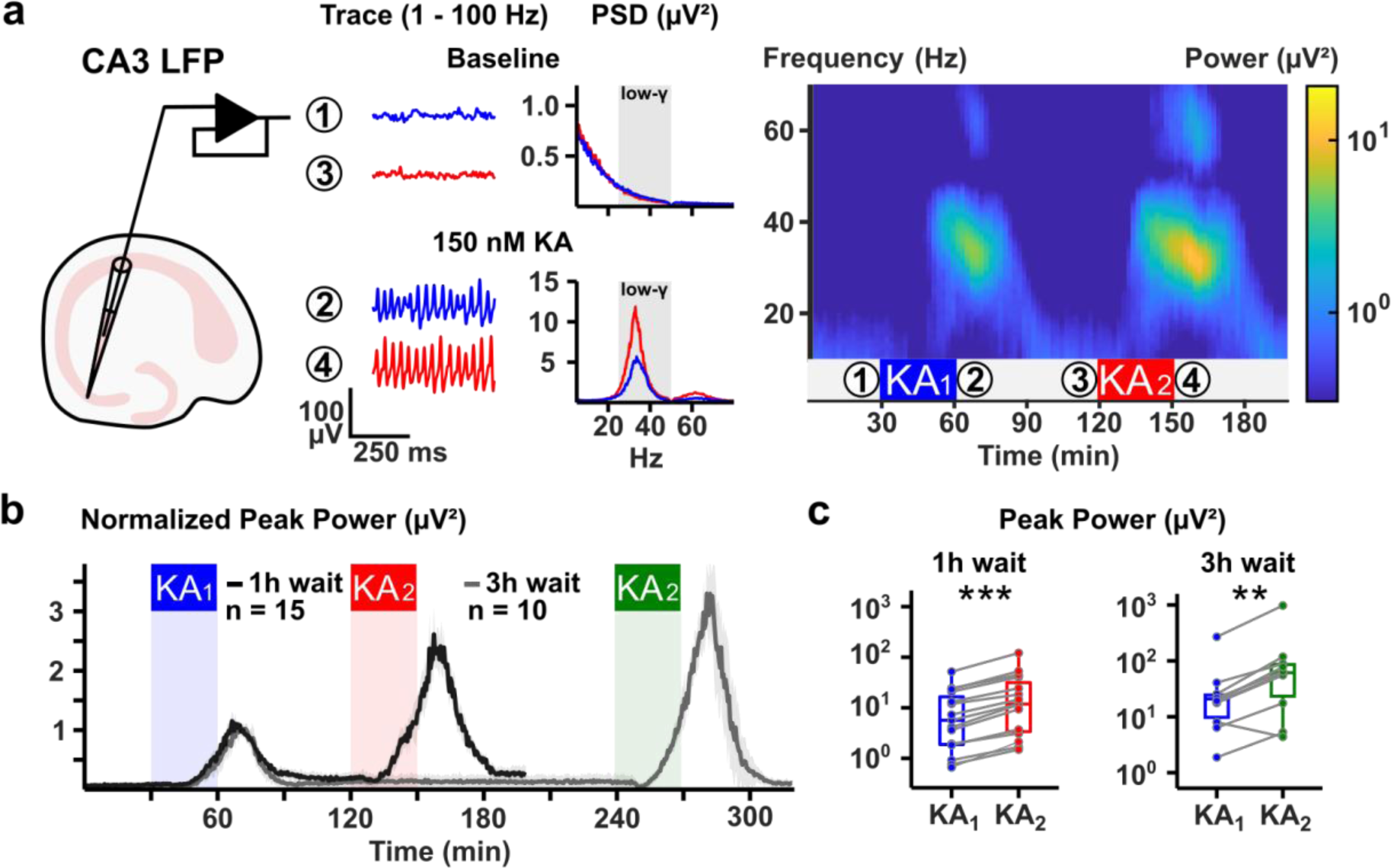
Gamma-potentiation in ex vivo mouse hippocampal CA3. **a** *Left*: Schematic of a hippocampal brain slice with an LFP electrode in the pyramidal cell layer of CA3. *Centre*: Close-up band-pass filtered traces (1 – 100 Hz) of the time periods preceding KA-application (‘Baseline’, insets 1 and 3 in the pseudocolor plot) and during maximum gamma-power in CA3 (‘150 nM KA-Gamma’, insets 2 and 4). The RMS-averaged power spectral density (PSD) was obtained over a 10-minute time window. Grey inset “low-gamma” in the PSDs denotes the window spanning 25 – 50 Hz. *Right*: Spectrogram of the entire recording. Blue and red insets ‘KA’ denote the time period of kainate application (KA, 150 nM). Colorbar on the right denotes the RMS-averaged power (log-scale). **b** Time-Power plot of peak power (15 – 49 Hz) for experiments with either 1- or 3-hour delay normalized to the first application period. Ribbons denote the 95% confidence interval by Tukey. **c** Paired boxplots of peak gamma-power in both application periods with either 1- or 3-hour delay. ** and *** denote p < 0.01 and < 0.001, respectively (Wilcoxon signed-rank test).

These findings suggest a ubiquitous plasticity rule, by which intrinsic gamma-activity in CA3 or CA1 induces long-term changes across the local microcircuit and enhances its oscillatory response on repeated identical stimulation. Our finding in transverse CA1 Mini-Slices, a slice model with reduced recurrent synaptic excitation onto pyramidal cells^30^, in particular points to plasticity acquired at local pyramidal cell-interneuron interactions.

### Calcium-permeable AMPA receptors essentially contribute to the gamma-rhythm, mediate and express gamma-potentiation

Gamma-oscillations arise from the recurrent interactions of synaptic excitation and inhibition provided by local pyramidal cells and interneurons, respectively^15^. Whereas GABAergic transmission is indispensable to oscillogenesis, we could analyze the contribution of individual glutamatergic components to gamma-synchronization and its potentiation using pharmacology. To specifically address synaptic excitation onto interneurons, we targeted calcium-permeable AMPA receptors (CP-AMPARs) with the open-channel blocker naphthyl-spermine (NASPM, 50 – 100 µM) and compared this to approaches targeting AMPARs globally (GYKI-53655, 50 µM) and/or NMDARs (D-AP5, 50 µM) (**Fig. 2A**).

**Figure 2.**
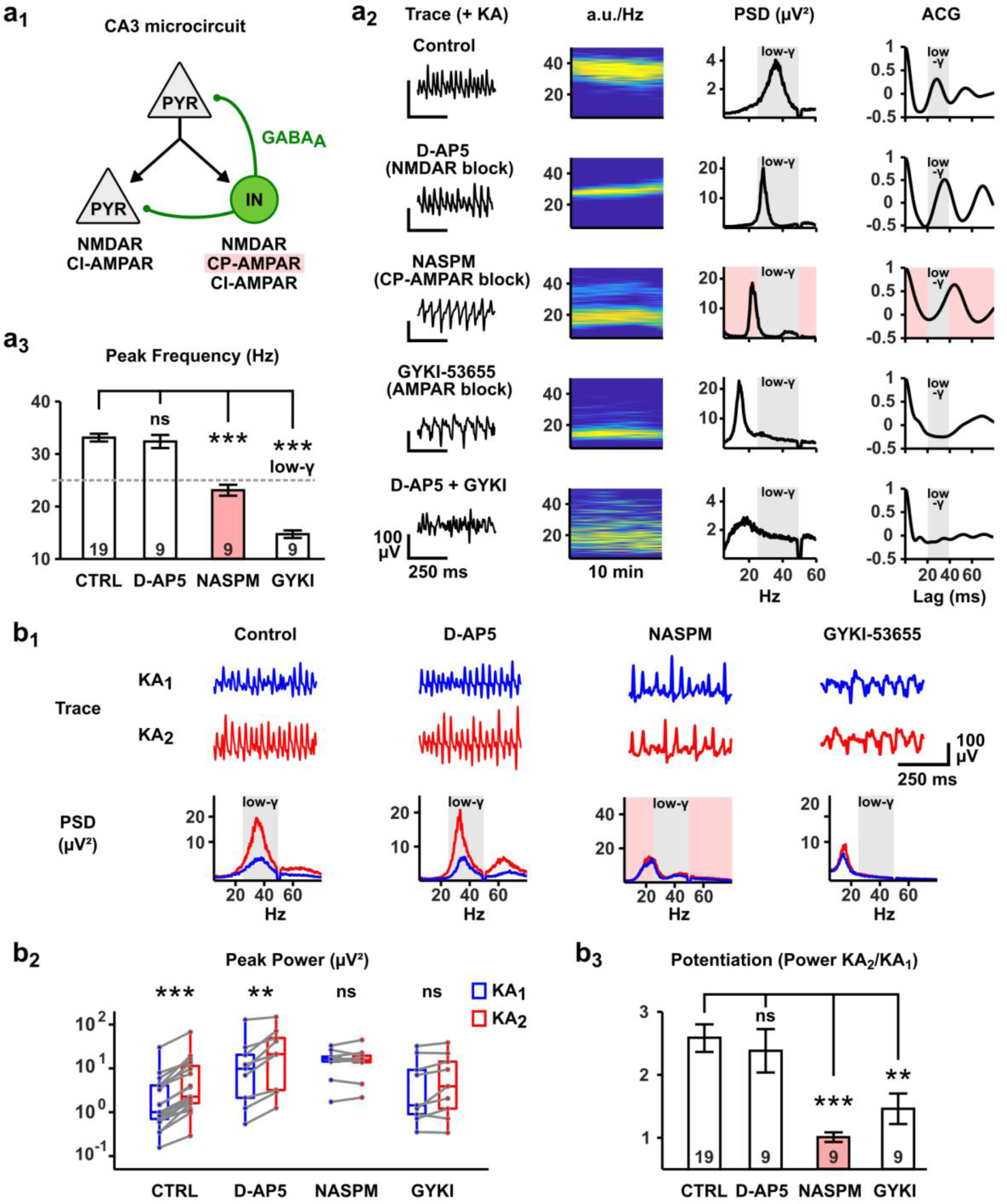
Calcium-Permeable AMPA receptors essentially contribute to the generation of the gamma-rhythm and mediate gamma-potentiation. **a1** Synapse-type specific targeting of glutamatergic transmission in CA3. In CA3, pyramidal cell (PYR) synapses targeting other PYRs express NMDA and calcium-impermeable AMPA receptors (CI-AMPARs). PYR synapses targeting interneurons (INs) additionally express calcium-permeable AMPA receptors (CP-AMPARs, pink inset). INs provide inhibition via GABAA-receptors. **a2** Contribution of ionotropic glutamate receptors to KA-induced oscillations. Traces, pseudocolor plots (power values plotted in arbitrary units, “a.u.”), PSDs and autocorrelograms (ACGs) of KA-induced oscillations after blockade of different ionotropic glutamate receptors. Grey insets “low-gamma” in the PSDs and ACGs denote the windows spanning 25 – 50 Hz and 20 – 40 ms, respectively. Pink inset highlights the specific blockade of CP-AMPARs by NASPM and reduction of peak frequency beneath the “low-gamma” range. There is no discernible oscillatory activity under co-application of D-AP5 and GYKI.

**a3** Barplot summarizing the peak frequencies in the oscillating conditions in **a2**. Grey dashed line indicates the lower border of the “low-gamma” frequency range (25 Hz). *** denotes p < 0.001 (Student’s t-test, corrected by Holm).

**b1** Exemplary potentiation experiments for the oscillating conditions in **a2**.

**b2** Paired boxplots of peak power values obtained in experiments shown in **b1**. ***, ** and ns denote p < 0.001, 0.01 and p > 0.05, respectively (Wilcoxon-Signed Rank test).

**b3** Barplot of the magnitude of gamma-potentiation for the experiments in **b1**. *** and ** denote p < 0.001 and p < 0.01, respectively (Student’s t-test, corrected by Holm).

CA3 LFP recordings revealed a fundamental contribution of CP-AMPARs to gamma-synchronization: When slices were pre-incubated with NASPM prior to stimulation with KA, the resulting rhythm was markedly slower than in controls (23.09 +/- 1.04 Hz, n = 9), yet clearly synchronous. This decelerating effect was even more pronounced after blocking all AMPARs with GYKI-53655 (14.71 +/- 0.72 Hz, n = 9). Finally, co-application of GYKI-53655 and D-AP5, which alone did not affect gamma-activity, all but prevented the emergence of oscillations in 8/9 slices tested (**Fig. 2A**). Therefore, synaptic excitation via AMPARs and NMDARs is itself paramount for KA-induced oscillations with a specific contribution of CP-AMPARs to reach and maintain peak frequencies in the low-gamma rhythm. This was confirmed in a set of pMEA experiments, permitting short-term drug application, in which gamma-activity was first established via application of KA, followed by an intermittent co-application of NASPM. Oscillations quickly de-synchronized when NASPM was applied, quantified as a variable reduction of either peak power and/or frequency, as well as a break-down of inter-site cross-correlation across CA3 recording sites, which partially recovered following washout of NASPM (**Supplementary Fig. 3A-C**). Concerning gamma-potentiation, when either GYKI-53655 or NASPM were pre-applied to our LFP protocol, the power of the resulting oscillations did not significantly increase during the second induction period (**Fig. 2B**, GYKI Power KA1: 1.43 [0.90 9.22] µV² vs. KA2: 3.89 [1.20 14.21] µV², n = 9, p = 0.10; NASPM Power KA1: 16.57 [14.27 18.52] µV² vs. KA2: 15.20 [13.79 19.52] µV², n = 9, p = 1, Wilcoxon Signed-Rank test). D-AP5, on the other hand, had no such effect, with potentiation remaining unchanged (Power KA2/KA1 Control: 2.58 +/- 0.21, n = 19 vs. D-AP5: 2.38 +/- 0.34, n = 9, p = 1, Student’s t-test). To exclude a possible contribution of KA receptors, we also tested the GluK1 antagonist UBP-302 (10 µM). Whereas pre-incubation with UBP-302 did raise the threshold for gamma-oscillation induction (400 nM instead of 150 nM), it did not prevent subsequent plasticity (**Supplementary Fig. 4**). Therefore, plasticity of KA-induced oscillations requires activation of AMPARs, specifically CP-AMPARs.

The dual role of CP-AMPARs in establishing gamma-activity and mediating subsequent plasticity limited the conclusions of our continuous LFP recordings. Combining our LFP protocol with subsequent pMEA recordings (6 × 10 grid, 100 µm inter-electrode distance) allowed us to observe gamma-oscillations at multiple sites covering CA3 and robustly compare independent samples. In multiple, parallel LFP recordings slices were first either treated with KA as previously or perfused with regular ACSF. After this initial period, all slices were left to rest for 1 – 3 hours and subsequently placed on pMEAs. Both treated and untreated slices were then stimulated with KA (200 nM, 3 minutes at 10 ml/min, 32 – 34° C), inducing low-gamma activity at sites covering the pyramidal cell layer of CA3c – CA3a (**Fig. 3A**). For each slice recorded, the electrode recording the highest gamma power was identified as the “lead electrode”, which was variably situated in CA3c, CA3b or CA3a. In both untreated (“KA1”) and treated (“KA2”) slices, peak power recorded at sites removed from the lead electrode gradually decreased as a function of intralaminar distance (100 µm to 1 mm, **Fig. 3A**). Comparing these distance-based values between slice conditions revealed that at all sites including the lead electrode, peak gamma power was markedly increased in treated slices compared to untreated slices (**Fig. 3A**). Therefore, gamma-potentiation is expressed in an activity-dependent manner and observed across the entire CA3 region.

**Figure 3.**
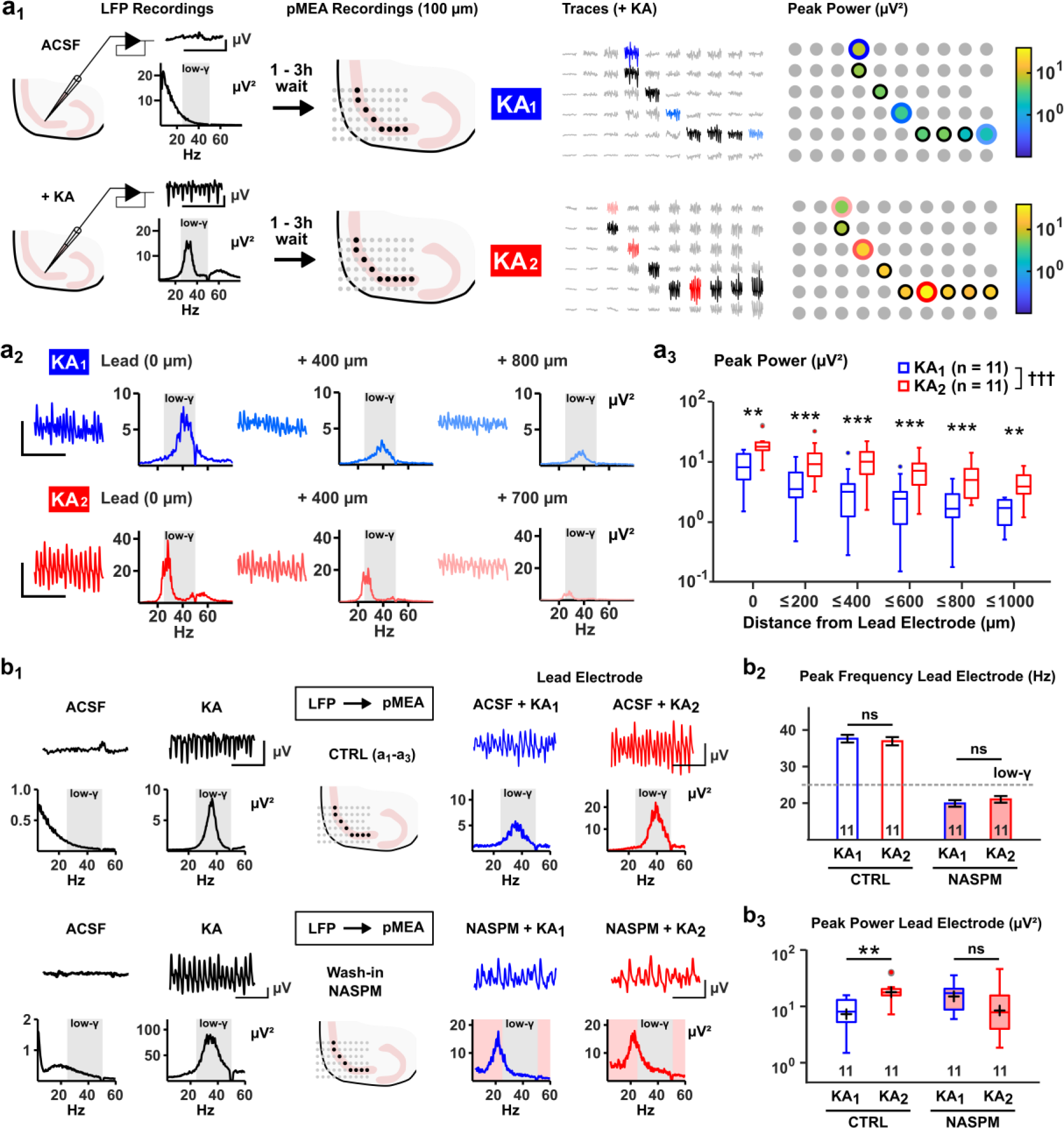
CP-AMPARs express gamma-potentiation in an activity-dependent manner across CA3. **a1** Combined LFP-pMEA protocol. *Left*, Slices are either left to rest (“Control”) or treated with KA, yielding low-gamma activity (“Gamma”). *Centre, Left*: After a waiting period slices are transferred to pMEA recordings with the selected electrodes (black) covering the pyramidal cell layer of CA3. *Centre, Right:* KA is applied to untreated (KA1, blue) and treated slices (KA2) and oscillations emerge. *Right* Heatmap of peak gamma-power over the selected electrodes for both slices. **a2** Exemplary traces and PSDs taken from the experiments in **a1**. Recordings from the electrodes with the highest power (“lead electrode”) are compared with those performed at distant sites, demonstrating a marked decrease of peak power over distance. **a3** Boxplot of peak power in both conditions pooled over the intralaminar distance from the lead electrode of each slice. **b1** CP-AMPARs express gamma-potentiation. Slices were grouped before LFP recordings and either treated with KA (30 minutes, 150 nM at 1.5 ml/min, 32-34° C) or left to rest before being transferred for pMEA recordings. Slices were then either maintained in regular ACSF (“Control”, same dataset as in **a-c**) or treated with NASPM (“NASPM-post”). The PSD of the “lead electrode” was calculated for analysis. **b2** Peak frequency of oscillations is reduced in pMEA “NASPM” conditions. **b3** In “Control” conditions, peak power of oscillations on the lead electrode is increased in pre-treated slices, but not when CP-AMPARs are blocked in the “NASPM-post” condition. ** and *** and ns denote p < 0.01 and < 0.001 and p > 0.05 for post-hoc Mann-Whitney U-tests corrected by Bonferroni-Holm. N is number of slices tested. Crosses denote p < 0.001 for Treatment (KA1, KA2) in a linear model (Peak Power ∼ Treatment + Intralaminar Distance).

We further made use of the combined LFP-pMEA approach to discern whether CP-AMPARs are required for just the induction of plasticity or also its subsequent expression. Slices were again divided into those stimulated with KA (thus, oscillating) and an unstimulated control group for initial LFP recordings and placed on pMEAs 1 – 3 hours after the first induction period. NASPM was then added to the bath before slices from both groups were again stimulated with KA (**Fig. 3B**). Peak frequencies of both unstimulated (NASPM + KA1) and stimulated (NASPM + KA2) slices were again decelerated (17 – 23 Hz) towards the control groups, yet comparably synchronous across multiple recording sites (**Supplementary Fig. 3D-E**). Other than in control recordings (Lead electrode Power KA1: 8.14 [5.33 13.16] µV² vs. KA2: 17.82 [15.84 20.46] µV², n = 11, p = 1.5 × 10^-3^, Mann-Whitney U-test), there was no difference of peak power at the lead electrode sites of unstimulated and stimulated slices in the after NASPM application (**Fig. 3B**, Lead electrode Power KA1: 17.34 [8.86 20.79] µV² vs. KA2: 7.93 [4.15 15.88] µV², n = 11, p = 0.15, Mann-Whitney U-test). Therefore, the increase of peak power is not just induced, but also expressed via the activation of CP-AMPARs.

In summary, we could pharmacologically dissect mechanisms of gamma-potentiation regarding the ionotropic glutamatergic transmission underlying its induction and expression: Whereas NMDARs and AMPARs contribute differentially to KA-driven hippocampal oscillations, the generation and subsequent potentiation of gamma-oscillations are mediated by AMPARs, expressed in an activity-dependent manner specifically by CP-AMPARs and independent of NMDARs and GluK1. This is strongly indicative of glutamatergic synaptic plasticity onto interneurons.

### A biophysically constrained microcircuit model of CA3 low-gamma oscillations predicts superior transfer of PVI-LTP to the resulting field potential

Gamma-oscillations induce glutamatergic long-term potentiation (LTP) onto both pyramidal cells (PYR) and predominantly CP-AMPAR expressing, parvalbumin-positive interneurons (PVIs)^27^. Whereas our pharmacological data suggested that an increase of CP-AMPA conductance at the PYR-PVI synapse underlies gamma-potentiation, LTP at the calcium-impermeable AMPAR (CI-AMPAR) expressing PYR-PYR synapse may, too, directly contribute to changes in the field potential or act heterosynaptically in recruiting PVIs. Lacking experimental tools to selectively target plasticity at PYR-PYR synapses, we turned to an *in silico* approach using computational modeling.

We developed a microcircuit model of CA3 low-gamma oscillations incorporating biophysically constrained, multi-compartmental PYRs (n=20) and PVIs (n=2) and recorded the extracellular “LFP” nearby the pyramidal cell somata. A detailed description of PYR and PVI cellular properties (PVIs adapted from^31^) as well as the model’s connectivity configuration can be found in **Methods** and **Supplementary Tables 1-3**. In this model, a baseline period of low-gamma activity can be reliably evoked by introducing an input population selectively targeting the PYR population for 1 second, mimicking our *ex vivo* approach using KA in a shorter time-scale (**Fig. 4A**, “Baseline” Power: 29.27 [26.70 30.42] µV²). We then simulated LTP at either CI-AMPAR containing PYR-PYR (PYR-LTP), CP-AMPAR containing PYR-PVI (PVI-LTP) synapses or both synapse types (PYR+PVI-LTP) by increasing their respective conductances by 50% and repeated the otherwise identical simulation (**Fig. 4B**). The PYR+PVI-LTP condition, which most closely approximated our *ex vivo* control experiments, reliably resulted in an increase of peak low-gamma power by a factor of 2.5 – 3 (**Fig. 4C-D**, “PYR+PVI-LTP” Power: 89.53 [83.69 98.01] µV², n = 10 trials). When simulations were repeated under identical conditions, yet applying just PYR-LTP, we still observed a significant increase of peak power, yet strongly reduced in magnitude (1.15-fold increase, **Fig. 4C-D**, “PYR-LTP” Power: 34.88 [28.94 36.53] µV², n = 10). In contrast, applying just PVI-LTP to our simulations sufficiently reproduced the increase of power seen in the PYR+PVI-LTP condition, in some cases outperforming it and revealing no additive interaction in combining PYR-LTP with PVI-LTP (**Fig. 4C-D**, “PVI-LTP” Power: 89.74 [82.70 93.21] µV², n = 10).

**Figure 4.**
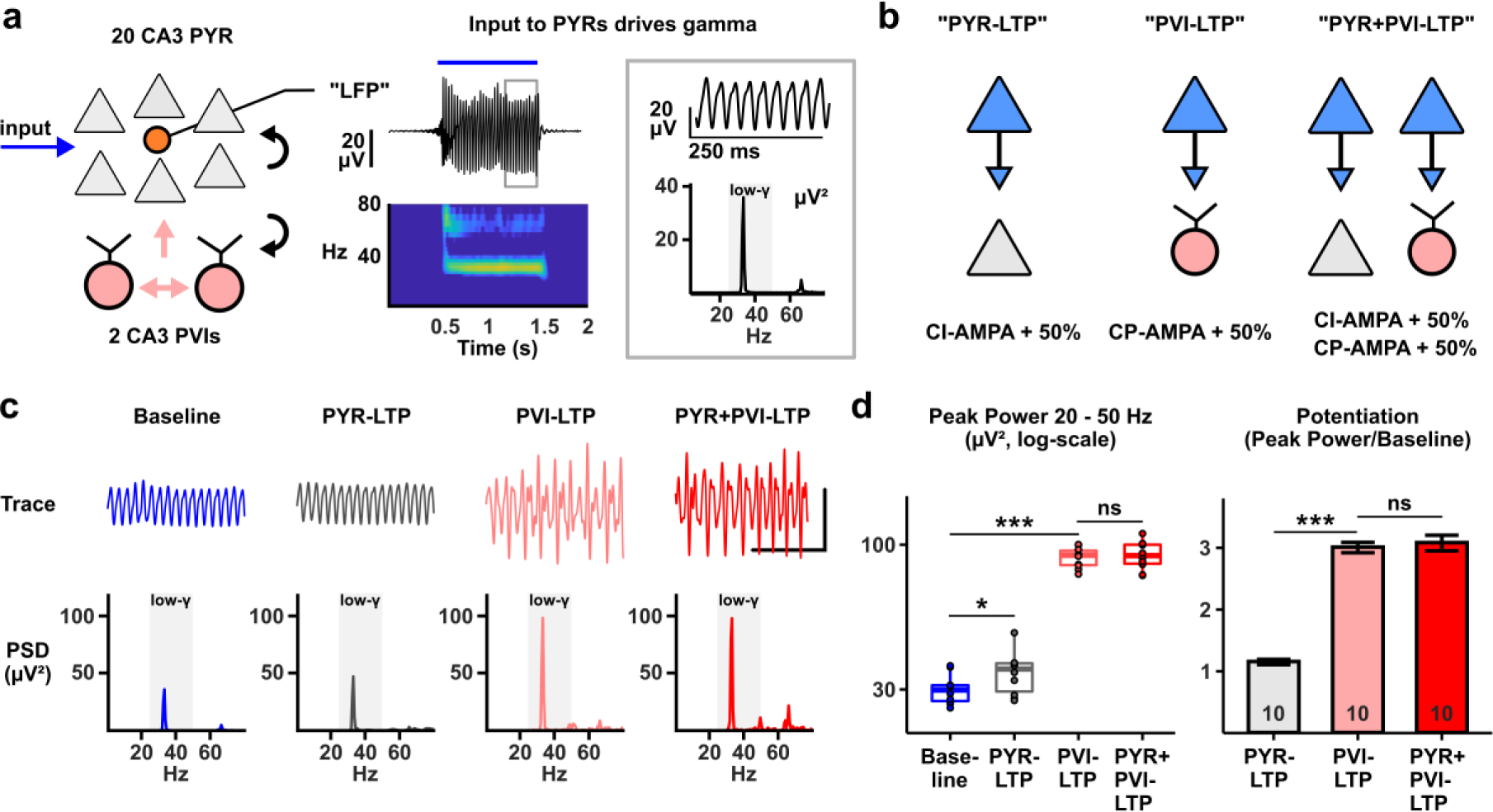
A biophysically constrained microcircuit model of CA3 low-gamma oscillations predicts superior transfer of PVI-LTP to the field potential. **a**, *Left*: Microcircuit model of CA3 low-gamma oscillations. A population of CA3 pyramidal cells (PYR, n = 20) is activated by an input population (blue arrow). The PYR population forms glutamatergic synapses (black arrows) onto both PYRs (calcium-impermeable AMPA [CI-AMPA] and NMDA conductances) and two CA3 PVIs (calcium-permeable [CP-AMPA] and NMDA conductances). The PVIs are connected to each other and form synapses onto the PYRs with GABAergic synapses (pink arrows). An *in silico* “LFP” electrode (orange) is positioned nearby the PYR somata. *Centre*: Exemplary trace of an LFP recording with pseudocolor plot below. A 1-second stimulation via the input population produces an oscillation between 30 – 40 Hz. *Right*: Close-up view of the trace and power spectrum of the entire 2-second recording. Grey inset marks the low-gamma frequency range (25 – 50 Hz). **b** *In silico* plasticity paradigms. Glutamatergic synapses (blue) formed onto PYRs (grey) and/or PVIs (pink) are modified to simulate cell-type specific plasticity. CI-AMPA conductances and/or calcium-CP-AMPA conductances are increased by 50% for the respective paradigm. **c** Traces and power spectra for one exemplary simulation. Inset denotes 50 µV by 250 ms. **d** Summary statistics including data from 10 random simulation trials per case. *Left*: Boxplots of peak low-gamma power for all conditions (* and *** denote p < 0.05 & p > 0.001, Wilcoxon-Signed Rank Test). *Right*: Barplot of the magnitude of potentiation (*** denotes p < 0.001, Student’s T-Test). Numbers in bars denote n = number of simulation trials.

Taken together, our simulations confirm a superior transfer of plasticity expressed at CP-AMPAR containing synapses, formed at PYR-PVI connections, to increasing low-gamma power. This is in line with our initial pharmacological data and predicts a substantial contribution of PVI-specific LTP to *ex vivo* gamma-potentiation.

### Mechanisms: Gamma-potentiation requires PV-specific mGluR5, mGluR1, PKC and PKA activation

PVI-LTP obtained during gamma-oscillations *ex vivo* can be prevented by unspecific concentrations (50 µM) of the group I mGluR antagonist MPEP^27^. In an initial set of LFP experiments, we tested the effect of MPEP on gamma-potentiation at a concentration specific to mGluR5 blockade (10 µM) as compared to co-application with D-AP5 and found that the magnitude of potentiation was robustly reduced by 50% in both cases (**Supplementary Fig. 5**). Due to a significant interaction of D-AP5 and MPEP in reducing the average peak frequency of oscillations (by roughly 2 Hz) and a modulating effect of both substances on peak gamma-power, all subsequent experiments involving MPEP were performed in the presence of D-AP5 (50 µM).

We next generated mice undergoing Cre/loxp-dependent postnatal ablation of mGluR5 under the PV-promoter^32^, enabling us to assess the cell-type specific contribution of mGluR5 to gamma-potentiation. Slices obtained from these animals (“PV-mGluR5 KO”) were compared to those obtained from littermates not expressing the loxp-mutation (“PV-mGluR5 WT”) and gamma-potentiation was quantified under application of MPEP (CTRL vs. MPEP, **Fig. 5A**). In both genotypes, gamma-oscillations were reliably induced with no apparent difference in peak gamma-power between control conditions (**Fig. 5A**, Power KA1 WT CTRL: 2.38 [0.41 4.27] µV², n = 16 vs. KO CTRL: 4.00 [1.20 34.57] µV², n = 28; p = 0.40, pairwise Mann-Whitney U-test). Regarding subsequent plasticity, MPEP again attenuated gamma-potentiation by approximately 50% compared to WT control slices (**Fig. 5A**, KA2/KA1 WT CTRL: 2.31 + /- 0.21, n = 16 vs. WT MPEP: 1.49 +/- 0.13, n = 19; p = 1.9 × 10^-3^, Student’s t-test), confirming our initial results. In slices from PV-mGluR5 KO animals, on the other hand, gamma-potentiation was already attenuated in control slices with no further reduction by MPEP (**Fig. 5A**, KA2/KA1 KO CTRL: 1.53 + /- 0.12, n = 28 vs. KO MPEP: 1.48 +/- 0.11, n = 19; p = 1.0, Student’s t-test). Indeed, gamma-potentiation in both KO conditions was similarly reduced towards WT control slices as via conventional blockade by MPEP (**Fig. 5A**). Moreover, the slightly attenuating effect of MPEP on peak gamma-frequency in the first induction period was not observed in KO slices (**Fig. 5A**). Therefore, effects of mGluR5 on both gamma-activity and subsequent gamma-potentiation are mediated specifically via its native expression on PVIs.

**Figure 5.**
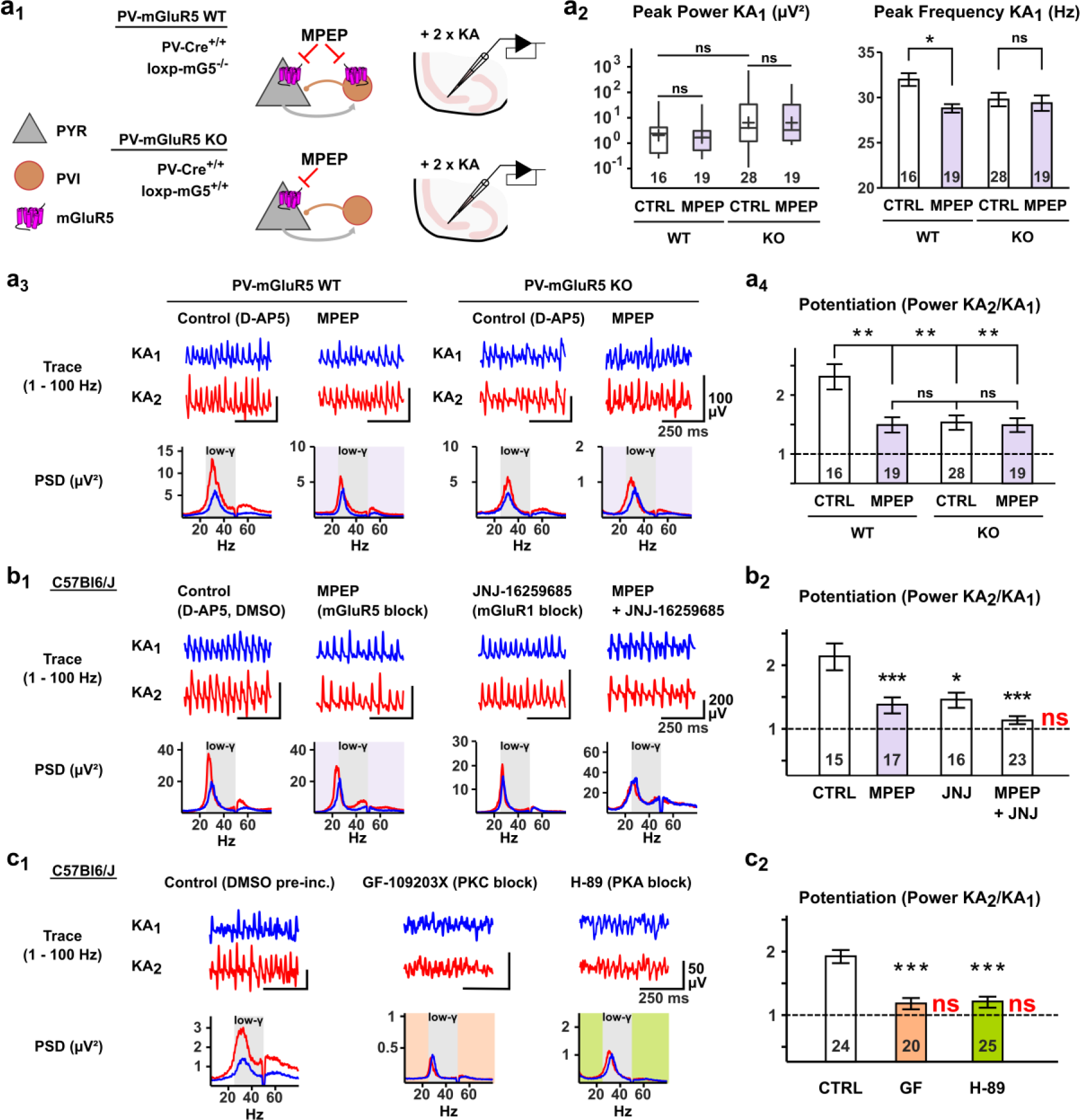
Gamma-potentiation requires PVI-specific mGluR5, mGluR1, PKC and PKA activation. **a1** Schematic of PV-mGluR5 experiments. Mice were bred homozygous for PV-Cre and either lacking (PV-mGluR5 WT) or homozygously expressing the loxp-mGluR5 mutation (PV-mGluR5 KO). MPEP blocks mGluR5 in a cell-type specific manner in PV-mGluR5 KO mice. Slices obtained from both animals were treated in our LFP paradigm. **a2** Baseline values of KA-induced gamma-oscillations in PV-mGluR5 WT and KO slices under application of MPEP. There is no difference in the peak power of oscillations between conditions. Application of MPEP reduces the peak frequency of oscillations in WT, but not KO animals. **a3** The attenuating effect of MPEP on gamma-potentiation requires mGluR5-expression in PVIs. *Above*: Exemplary traces of Control and MPEP conditions for PV-mGluR5 and PV-mGluR5 KO slices during the first (KA1) and second (KA2) induction period. *Below*: Corresponding PSDs for each condition. All experiments performed in 50 µM D-AP5. **a4** Barplot of the magnitude of gamma-potentiation for the experiments in **a3**. **b1** Gamma-potentiation requires group I mGluRs, PKC and PKA. *Above*: Exemplary traces of Control, MPEP, JNJ and MPEP + JNJ conditions during the first (KA1) and second (KA2) induction period with corresponding PSDs for each condition beneath. Experiments were performed in the presence of 50 µM D-AP5 and 0.01% DMSO. **b2** Barplot of the magnitude of gamma-potentiation for the experiments in **b1**. **c1** Gamma-potentiation requires PKC and PKA. Traces and PSDs as in **b1.** All slices were pre-incubated for 1 hour in 0.01% DMSO before experiments. **c2** Barplot of the magnitude of gamma-potentiation for the experiments in **c1**. Numbers in plots denote the number of slices tested per condition. ***, ** and ns denote p < 0.001, p < 0.01 and p > 0.05 (Student’s t-test corrected by Bonferroni-Holm in barplots, Mann-Whitney U-test or Wilcoxon Signed-Rank in boxplots). Red insets “ns” in potentiation plots mark groups with no significant increase of peak gamma-power in the second induction period (Wilcoxon Signed-Rank test). Purple, orange and green insets mark the application of MPEP, GF 109203X and H-89, respectively.

mGluR1-coactivation may account for the remaining 50% of gamma-potentiation, as both group I mGluRs similarly contribute to CP-AMPAR mediated PVI-LTP by entraining protein kinase C (PKC)^33, 34^. Beyond such a canonical Gq-pathway, protein kinase A (PKA) or activation of voltage-gated calcium channels (VGCCs), too, contribute to a diverse set of plasticity mechanisms putatively relevant to our paradigm. We tested these assumptions in a set of pharmacological LFP experiments in wild-type mice (**Fig. 5B**). First, pre-application of the mGluR1 antagonist JNJ-16259685 (0.3 µM) reduced the magnitude of gamma-potentiation towards control conditions (D-AP5, DMSO) by roughly 50% (**Fig. 5B**, KA2/KA1 CTRL: 2.13 + /- 0.20, n = 15 vs. JNJ: 1.45 +/- 0.11, n = 16; p = 2 × 10^-3^, Student’s t-test) and slightly reduced the peak frequency of oscillations (data not shown), comparable to sole application of MPEP (KA2/KA1 MPEP: 1.36 + /- 0.12, n = 17, p = 7 × 10^-4^ vs. Control). When both substances were co-applied, we found no significant gamma-potentiation during the second induction period (**Fig. 5B**, MPEP + JNJ Peak Power KA1: 34.16 [11.91 67.14] µV² vs. KA2: 35.83 [15.90 77.57] µV², n = 23, p = 0.23, Wilcoxon Signed-Rank test), confirming the additive requirement of group I mGluR activation in our protocol. In a second set of experiments, slices were first pre-incubated for 1 hour with either the PKC antagonist GF 109203X (3 µM), the PKA antagonist H-89 (3 µM) or a respective DMSO control. Gamma-oscillations could be reliably induced in all conditions, whereas oscillations induced following blockade of PKC, but not PKA, displayed a decreased initial peak gamma-power (data not shown). However, both pre-incubation with GF 109203X or H-89 entirely prevented gamma-potentiation during the second induction period, demonstrating a requirement of both PKC and PKA activation for gamma potentiation (**Fig. 5C**, Peak Power Control KA1: 1.46 [0.77 6.18] µV² vs. KA2: 3.22 [1.64 9.98] µV², n = 24, p = 1.2 × 10^-7^; GF KA1: 0.30 [0.14 0.85] µV² vs. KA2: 0.41 [0.15 1.02] µV², n = 20, p = 0.18; H-89 KA1: 1.40 [0.30 5.91] µV² vs. KA2: 1.58 [0.46 2.68] µV², n = 25, p = 0.62, Wilcoxon Signed-Rank test). Finally, when either the L-Type VGCC antagonist Nifedipine (10 µM) or the T-Type antagonist ML-218 (5 µM) were pre-applied, neither the induction of oscillations nor their subsequent potentiation was significantly affected (**Supplementary Fig. 6**).

This completes a mechanistic profile of gamma-potentiation nearly identical to known plasticity rules of glutamatergic LTP onto PVIs: Plasticity of gamma-power is independent of NMDARs and VGCCs and instead requires CP-AMPARs, group I mGluRs and PKC^34^ with an additional requirement of PKA activation. This profile is directly tied to PVIs by the requirement of the cell-type specific expression of mGluR5.

### DREADD-based metabotropic manipulation of PVIs determines the induction of gamma-potentiation

Whereas our data from PV-mGluR5 KO slices accounted for the pharmacological effect of MPEP (50% of overall potentiation), open questions remained regarding the residual mGluR1 component and the locus of PKA action. We addressed this by applying two DREADD-based strategies specific to PVIs and bred mice expressing either hM4Di^35^ or HA-tagged hM3Dq under the PV-promoter alongside the fluorescent marker Ai9 (“PV-Ai9-hM4Di” and “PV-Ai9-hM3Dq” animals, respectively). In slices obtained from these animals, the respective DREADD-pathway can be reliably activated with the selective compound deschloroclozapine (DCZ)^36^.

Slices obtained from PV-Ai9-hM4Di animals were pre-incubated with DCZ (3 µM) prior to KA-application, arguably reducing intracellular cAMP levels and downstream thereof PKA activity^37^ (**Fig. 6A**). To preclude non-specific effects of DCZ, concomitant experiments were performed in mice lacking hM4Di-expression (“PV-Ai9”). Neither in slices from PV-Ai9 nor PV-Ai9-hM4Di animals did DCZ affect the induction or maintenance of gamma-activity during the first induction period in LFP recordings (data not shown). However, in slices from PV-Ai9-hM4Di animals, subsequent gamma-potentiation was completely prevented by DCZ (**Fig. 6B**, Power KA2/KA1 PV-Ai9-hM4Di Control: 2.37 +/- 0.23, n = 15 vs. DCZ: 1.09 +/- 0.10, n = 15, p = 2.6 × 10^-5^), but not in slices from PV-Ai9 animals (PV-Ai9 Control: 2.24 +/- 0.23, n = 13 vs. DCZ: 2.25 +/- 0.19, n = 13, p = 0.96, Student’s t-test), indicating that the effect of PKA we had observed in wild-type animals can be attributed to its specific activation in PVIs.

**Figure 6.**
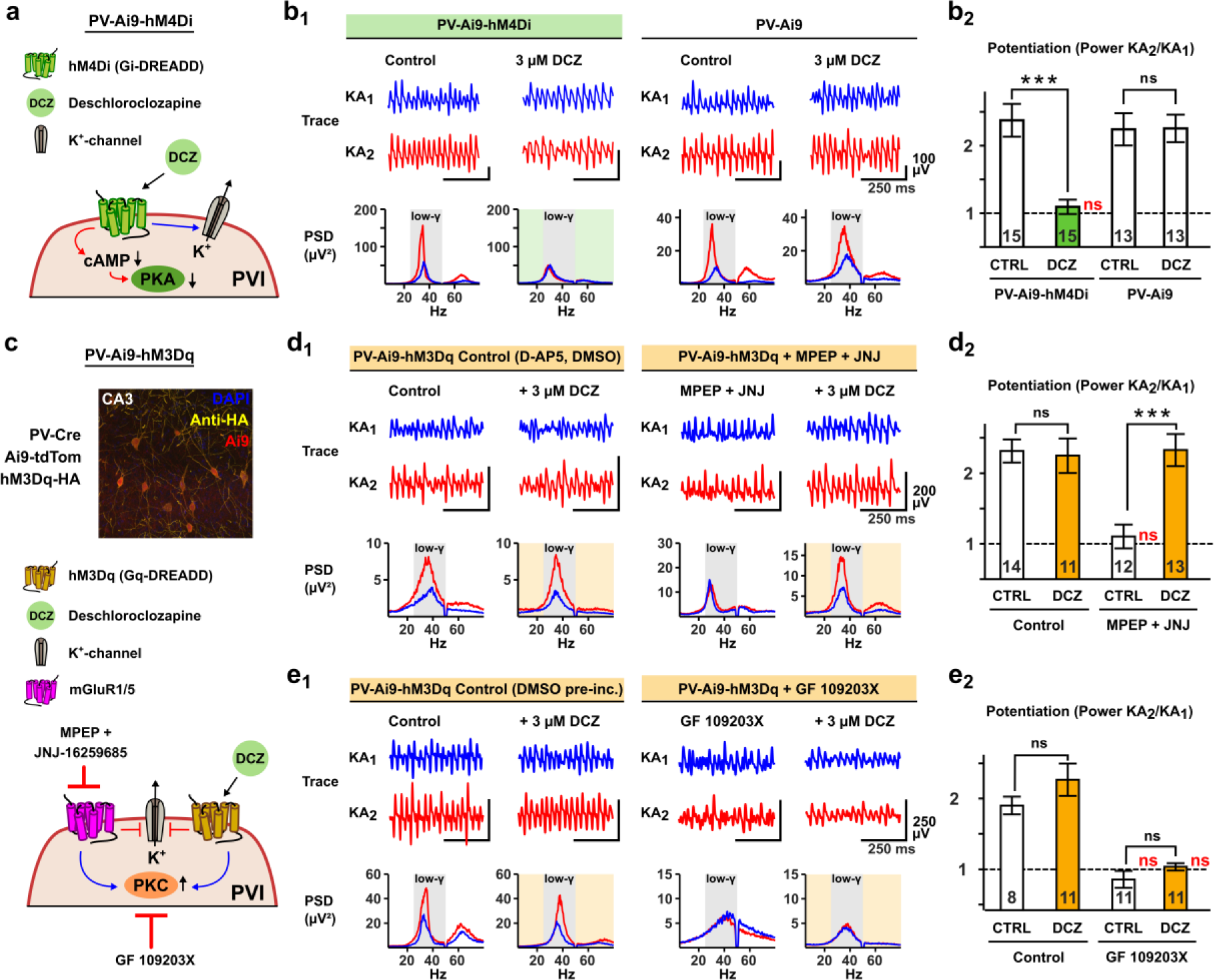
PVI-specific DREADD manipulations determine the induction of gamma-potentiation. **a** Schematic of PV-Ai9-hM4Di model. By activating hM4Di in PVIs with DCZ, both intracellular activity of cAMP and PKA are reduced in PVIs and cells are putatively hyperpolarized via potassium channel activation. **b1** DCZ prevents gamma-potentiation in the PV-Ai9-hM4Di model. *Left:* Exemplary traces of Control and DCZ conditions for both genotypes during the first (KA1) and second (KA2) induction period. *Below:* Corresponding PSDs for each condition. **b2** Barplot of potentiation in each condition. **c** *Above* HA-tagged hM3Dq is broadly expressed in CA3 PVIs (30x magnification). *Below* Schematic of PV-Ai9-hM3Dq model and pharmacological approach. By activating hM3Dq in PVIs with DCZ, potassium channels are closed, depolarizing the cells, and PKC is activated. mGluR1 and mGluR5 target the same pathways, which is prevented with MPEP and JNJ-16259685. PKC is antagonized with GF 109203X. **d1** DCZ rescues gamma-potentiation in PV-Ai9-hM3Dq animals after blockade of group I mGluRs. *Above:* Exemplary traces of Control and DCZ conditions for both the regular paradigm and under application of MPEP and JNJ during the first (KA1) and second (KA2) induction period. *Below:* Corresponding PSDs for each condition. All experiments performed in 50 µM D-AP5 and 0.01% DMSO. **d2** Barplot of potentiation in each condition. **e1** The DCZ-dependent rescue is prevented by blockade of PKC. *Above:* Exemplary traces of Control and DCZ conditions for both the regular paradigm and under application of GF 109203X during the first (KA1) and second (KA2) induction period. *Below:* Corresponding PSDs for each condition. All experiments performed after pre-incubation with 0.01% DMSO for one hour. **e2** Barplot of potentiation in each condition. Coloured insets in PSDs and barplots highlight the application of DCZ in the respective condition. *** and ns denote p < 0.001 and p > 0.05 (Student’s t-test corrected by Bonferroni-Holm. Red insets “ns” in potentiation plots mark groups with no significant increase of peak gamma-power in the second induction period (Wilcoxon Signed-Rank test).

In slices from PV-Ai9-hM3Dq animals, we applied the inverse strategy: If gamma-potentiation is prevented by blockade of endogenous Gq-receptors or PKC, activation of the xeno-receptor hM3Dq in PVIs may be sufficient to rescue blockade of the mGluRs, yet not blockade of downstream PKC (**Fig. 6C**). On the other hand, hM3Dq-activation and subsequent depolarization of PVIs independent of PKC may itself drive PVI firing and thus induce gamma-activity, as has been shown for optogenetic strategies^8^. We tested these assumptions in multiple steps: First, when applied by itself, DCZ was insufficient to induce any form of network synchronization, highlighting the requirement of synaptic excitation we had demonstrated earlier (**Supplementary Fig. 7**). Second, when mGluR5 and mGluR1 were blocked with MPEP (10 µM) and JNJ-16259685 (0.3 µM), gamma-potentiation was again prevented in slices from PV-Ai9-hM3Dq animals (Power KA2/KA1 PV-Ai9-hM3Dq Control: 2.31 +/- 0.15, n = 14 vs. MPEP + JNJ: 1.10 +/- 0.16, n = 12, p = 7 × 10^-5^). This was entirely rescued when DCZ was co-applied with KA, with the magnitude of potentiation nearly identical to control experiments (**Fig. 6D**, Power KA2/KA1 PV-Ai9-hM3Dq MPEP + JNJ + DCZ: 2.32 +/- 0.21, n = 13). Third, gamma-potentiation was similarly prevented by pre-incubation with GF 109203X (3 µM, DMSO). In this case, co-application of DCZ failed to reinstate gamma-potentiation (**Fig. 6E**, Power KA2/KA1 PV-Ai9-hM3Dq Control: 1.93 +/- 0.12, n = 8 vs. GF: 0.86 +/- 0.11, n = 11 & GF + DCZ: 1.04 +/- 0.05, n = 11). Therefore, gamma-potentiation can be entirely accounted for by metabotropic signaling in PVIs dependent on a Gq/PKC- and a Gi-sensitive PKA-pathway, with DREADD-based strategies in PVIs functioning as effective on/off-switches for network plasticity.

## Discussion

Studying the intersections of interneuron plasticity, oscillatory activity and learning has emerged as a challenging, yet promising field of research^38, 39^. Particularly in the hippocampus the learning of context has been associated with changes of oscillation patterns: Following contextual fear conditioning *in vivo*, oscillations in the theta^20^, gamma^22^ and ripple^20, 21^ frequency bands undergo plastic changes in CA1. In CA3, theta-gamma coupling increases during successive trials of item-context association^4^ and low-gamma power increases following object learning, a change associated with improved assembly formation of CA3 pyramidal cells^6^. Whereas targeted cell-type specific manipulations have linked some of these phenomena to the activation of interneurons, an understanding of cause and effect of oscillatory plasticity remains elusive. We previously demonstrated in a first step that hippocampal gamma activity itself induces plasticity of sharp-wave ripple complexes *in vivo* and *ex vivo*, which was associated with synaptic plasticity on both pyramidal cells and interneurons^27^.

Here, we demonstrate a general plasticity mechanism intrinsic to hippocampal network oscillations by which gamma power is increased upon repeated exposure to an equal excitatory stimulus (**Fig. 1**). This plasticity mechanism – gamma potentiation – is embedded in a reciprocal relationship with PVI synaptic plasticity (**Figs. 4-6**): Evoking network activity in the gamma-frequency range induces long-lasting plasticity onto PVIs^27, 33, 34, 40^, which in turn translates to subsequent network activity as a function of peak gamma-power. Besides our PVI-specific interventions (**Figs. 5-6**), this is supported by two inductive approaches: *Ex vivo*, our data from CA1-Mini slices (**Supplementary Fig. 2**), a slice model of reduced recurrent excitation on pyramidal cells^30^, demonstrates that local pyramidal cell–interneuron interactions are sufficient for the induction of oscillation plasticity. *In silico*, our micro-scale model of CA3 connectivity containing both PYR-PYR and PYR-PVI synapses further corroborates that plasticity at the PYR-PVI synapse critically outperforms plasticity at the PYR-PYR synapse in increasing low-gamma power (**Fig. 4**). However, we do not exclude possible additional contributions of other interneuron sub-types: Somatostatin-positive interneurons, which to an extent co-express PV, also contribute to gamma-oscillations in the hippocampus^41^ and neocortex^42, 43^ and may similarly contribute to their plasticity. Yet within this study, plasticity at the CA3 pyramidal cell-PVI synapse is both sufficient and mandatory to modulate the gamma rhythm.

We investigate this relationship mechanistically using a robust (**Supplementary Fig. 1**) *ex vivo* paradigm, permitting the analysis of gamma-activation states under maximal experimental control and circumventing confounders of *in vivo* oscillation power such as running speed, respiration rate or cross-frequency coupling^44, 45^ as well as spatial limitations concerning the origin^5, 6, 46^ and focality^47^ of oscillations. In the approach used here, gamma-activity is gradually introduced to the isolated hippocampal network and tapered under constant control of external excitation via defined application of kainate. This allows the precise mechanistic dissection of plasticity rules with pharmacological and genetic tools. Plasticity in our protocol occurs in an activity-dependent manner and already has effect after short-to intermediate term delays in subsequent episodes (1 – 3 hours), predating alterations in transcriptional and anatomical PVI properties^24, 25^ and making changes of transmembrane conductances the most likely mechanism. Translating these *ex vivo* conclusions to the ground-truth *in vivo* gamma-rhythm will require temporally precise synapse-specific tools^48^ and an experimental environment closely controlling the emergence and re-instatement of gamma-oscillations on short time scales.

The specific synaptic recruitment of PVIs via CP-AMPARs is the decisive determinant in procuring and expressing gamma-potentiation, as demonstrated by both our *ex vivo* and *in silico* data sets (**Figs. 2-4**). Through their rapid activation and decay kinetics, CP-AMPARs enable the temporally precise integration of converging inputs on and co-activation of PVIs^49, 50^, which our data suggests is mandatory for the induction of >30 Hz oscillations themselves. Therefore, increased CP-AMPAR conductance is a convincing mechanistic candidate underlying gamma potentiation. Given our previous findings for sharp-wave ripple associated plasticity^27^ and present modelling predictions, this points to the induction of synaptic glutamatergic LTP onto PVIs (PVI-LTP) as the primary facilitator of gamma-potentiation. Importantly, the specific requirement of CP-AMPAR activation reported here separates our protocol from previous optogenetic or chemogenetic manipulations of network plasticity purportedly relying on the isolated activation of PVIs, but which cannot exclude coincident glutamatergic drive^20, 21, 51^. Whether downstream effectors of PVI-LTP such as intrinsic membrane excitability^52^ and/or an increased transmission at GABAergic output synapses similarly contribute to the resulting field potential remains to be determined^53–55^.

Like PVI-LTP in the dentate gyrus and gamma-mediated PVI-LTP in CA3^27, 34^, gamma-potentiation is independent of NMDAR activation and instead requires group I mGluRs and downstream PKC, in line with a canonical Gq-cascade. We tie this to PVIs with two genetic strategies: The conditional ablation of mGluR5 in PVIs as well as PV-specific Gq-DREADD activation. Moreover, both our pharmacological and chemogenetic evidence using hM4Di-activation suggest the requirement of a second pathway recruiting PKA (**Figs. 5-6**). A dual requirement of two kinase pathways may act as a gating mechanism for network plasticity^56^: If gamma-oscillations were both induced and amplified via CP-AMPAR activation alone, local networks would be exposed to run-off dynamics in which oscillation power continuously increases. Expanding such a model by a pre-requisite of metabotropic co-stimulation stabilizes network dynamics and discriminates between sub- and supra-critical stimuli to the hippocampal network during learning. This mirrors a recent proposal of three-factor plasticity in interneurons^39^, in which mGluRs function as effective detectors of converging activity during oscillations^57^, and putative activators of PKA (e.g. dopamine, noradrenaline) encode novelty^58, 59^. Whether the Gq/PKC- and PKA-pathways effectuate plasticity independently, converge with each other or interact via cross-talk^60^ remains beyond the scope of this study yet may provide crucial insights into how targeting PVI-LTP translates into network oscillations.

The mechanistic composition of gamma-potentiation provides a promising framework for the understanding and potential treatment of neuropsychiatric diseases associated with aberrant network oscillations. The PV-mGluR5 knockout model used here (**Fig. 5**) displays core features of neurodevelopmental disease^32^, establishing a direct link between insufficient *ex vivo* gamma-plasticity and disrupted hippocampal behaviour. mGluR5 activity is downregulated in post-mortem tissue of schizophrenia patients^61^ and, conversely, positive allosteric modulation of mGluR5 has been successfully targeted in animal models of schizophrenia^62^, a disease etiologically tied to PVI dysfunction and disrupted network oscillations. The link between PVI plasticity, gamma-potentiation and disease is further supported by a string of recent pre-clinical studies successfully treating symptoms of neurodevelopmental disorders: Similar to the bimodal control of PVI-dependent DREADD manipulations over gamma potentiation (**Fig. 6**), hM4Di-inhibition of PVIs induces deficits of cognitive performance^21, 63^, while PV-specific hM3Dq-activation rescues such deficits in animal models of schizophrenia as well as disruptions in network oscillations^63–65^. Further, the physiological induction of gamma-oscillations via sensory stimuli (gamma entrainment using sensory stimuli, GENUS) is itself effective in treating animals models of Alzheimer’s disease^66^ and schizophrenia^67^, whereas the exact mechanism underlying these treatments has been recently contested^68^. An inert, bidirectional cortical mechanism of gamma-rhythm plasticity via synaptic plasticity of PVIs may therefore lie at the core of future clinical interventions and serve to inform therapeutic strategies.

## Materials and Methods

### Animals

Adolescent male C57Bl6/J mice (p45 – p70) were used for experiments in wild-type animals. For experiments involving cell-type specific manipulations of PVIs, PV-Cre animals (The Jackson Laboratory, Stock No. 017320) were crossbred with either Ai9 (Jackson, Stock No. 007909), loxp-mG5 (see^32^, kindly provided by Peer Wulff), Flex-hM4Di (see^35^, kindly provided by Benjamin Rost) or loxp-hM3Dq (The Jackson Laboratory, Stock No. 026220) animals and experiments performed from adolescent (p45 – p70) offspring of both sexes. Animal procedures were conducted in accordance with the guidelines of the European Communities Council and the institutional guidelines approved by the Berlin Animal Ethics Committee (Landesamt für Gesundheit und Soziales Berlin, T0045/15 and T-CH0014/23). All efforts were made to minimize animal suffering and to reduce the number of animals used.

### Slice Preparation

Mice were deeply anaesthetized with isoflurane and decapitated. Their brains were removed and immersed in ice-cold sucrose solution (in mM: Sucrose 75, NaCl, 87; KCl, 2.5; NaHCO3, 25; NaH2PO4, 1.25; MgCl2, 3; CaCl2, 0.5; glucose, 10) saturated with carbogen gas (95% O2 / 5% CO2). The brain was cut into 400 µm thick horizontal slices containing the hippocampal formation with a vibratome (Leica VT 1200S, Leica Biosystems, Germany) and slices were transferred to interface-type recording chambers perfused with warm and carbogenated artificial cerebrospinal fluid (ACSF; in mM: NaCl, 129; KCl, 3; NaHCO3, 21; NaH2PO4, 1.25; MgSO4, 1.8; CaCl2, 1.6; glucose, 10; 32 – 34° C; flow rate: 1.5 ml/min) and were left to incubate for 2 h before recordings. To obtain CA1-Mini slices, slices were cut in the interface chamber shortly after preparation with a surgical blade to separate CA1 from CA2 and the subiculum under consideration of CA1 dendrite morphology.

For experiments involving pharmacological agents, drugs were added to ACSF at least one hour prior to recording. In experiments involving GF109203X or H-89, slices were left to recover post-slicing in a submerged-type sucrose beaker at 34° C for one hour containing the respective substance at its final concentration before being transferred to interface chambers. Corresponding control experiments were performed in the presence of 0.01% DMSO in the beaker.

### Electrophysiology

Local field potentials were recorded from stratum pyramidale of hippocampal CA3 (and/or CA1 when indicated) with glass pipettes filled with ACSF (1 - 10 MΩ). Recordings were amplified by EXB-EXT-02B amplifiers (npi Electronic, Germany), low-pass filtered at 1 kHz, sampled at 5 kHz by a CED 1401 AD-converter (Cambridge Electronic Design [CED], UK) and saved to disk via Spike2 software (CED, UK). Following an initial 30-minute period of recording baseline activity, network oscillations were induced in one or two separate periods by bath application of 150 - 400 nM kainate (KA) over a period of 30 minutes with a 60 minute “resting” interval between both application periods. In a subset of experiments the resting period was extended to three hours. After the second application of KA, slices rested for up to one hour before terminating the recording. Pipette resistances were monitored before and after recordings with an EXB-REL08B electrode resistance meter (npi) and recordings discarded if resistances deviated > 10%.

Perforated multi-alectrode array (pMEA) recordings were performed on a MEA2100-HS(2x)60 system (Multichannel Systems) using 60pMEA100/30iR-Ti pMEAs (Multichannel Systems). Slices were transferred from their previous interface storage to the pMEA and carefully placed above the electrodes with the aim of maximal coverage of the pyramidal cell layer (identified visually under magnification and confirmed by the positive polarity of spontaneous sharp-wave ripple complexes). Slices were kept in place via a continuous negative pressure supplied by a constant vacuum pump (CVP-230V, Multichannel Systems) and allowed to rest in position for 15 – 20 minutes before recording. After a brief recording of baseline activity confirming the absence of ambient oscillations, oscillations were induced via bath-application of 200 nM KA (10 ml/min flow rate, 32 – 34° C). The dual headstage configuration of the MEA2100-HS(2x)60 system allowed us to test individual treated and untreated slices in time control.

### Drugs

Kainate (Tocris, KA), D-AP5 (Cayman Chemicals), MPEP (Cayman Chemicals), JNJ-16259685 (Tocris, JNJ), GYKI-53655 (hellobio), Naphthyl-spermine (Cayman Chemicals, NASPM), GF109203X (Tocris, GF), H-89 (Cayman Chemicals), UBP-302 (Tocris, UBP), Deschloroclozapine-dihydrochloride (Cayman Chemicals), Nifedipine (Tocris) and ML-218 (Tocris) and were dissolved in either de-ionized water or DMSO (JNJ, GF, H-89, UBP, VU, Nifedipine, ML-218) and stored in aliquots at -20° C. Aliquots were dissolved in ACSF immediately prior to the experiments. Experiments involving Nifedipine were conducted in the dark.

### Computational modeling

#### Model implementation and availability

We conducted all modeling simulations using the NEURON (NEURON v7.6) Simulator (Hines & Carnevale, 1997) on a High-Performance Computing Cluster (HPCC) with 111 CPU cores, running on a 64-bit CentOS Linux operating system. The source code and datasets used to generate **Fig. 4** are publicly available; please refer to the Code and Data Availability section for the link.

### Model neurons

#### PV Basket Cell Model

The multi-compartmental models of the CA3 PV Basket cells (PVIs) (n=2) used in this study were adapted from previously published work^31^. These models have been extensively validated against experimental data and have been shown to accurately capture the intrinsic, active, and morphological properties of PVIs (for more detailed information, please refer to the aforementioned publication).

#### Pyramidal Neuron Model

The CA3 pyramidal neuron cell model (PYR) was simulated based on the Hodgkin-Huxley formalism and consists of six compartments: one soma and five dendrites. The model simulates one proximal and two distal apical dendrites, as well as two basal dendrites. It includes a Ca^2+^ pump and buffering mechanism, Ca^2+^ activated slow AHP and medium AHP potassium (K^+^) currents, an HVA L-type calcium (Ca^2+^) current, an HVA R-type Ca^2+^ current, an LVA T-type Ca^2+^ current, an h current, a fast sodium (Na^+^) current, a delayed rectifier K^+^ current, a slowly inactivating K^+^ M-type current, and a fast inactivating K^+^ A-type current. The current mechanisms were distributed in a non-uniform way along the somatodendritic compartments (detailed information about the passive and active properties of the PYR model, including conductance values, please see **Supplementary Table 1**). The active and passive properties of the PYR model were validated against *in vitro* experimental data from CA3 recordings^69^. This was done to ensure that the *in silico* model reproduces the electrophysiological profile of the *in vitro* CA3 pyramidal cells (for more detailed information, please see **Supplementary Table 2**).

#### Synaptic Properties

The PYR models were equipped with CI-AMPA, NMDA, and GABAA synapses, while the PV BC models had CPAMPA, NMDA, GABAA, and autaptic GABAA synapses. The synaptic properties were validated against previously published data^70–72^. The conductance values for each synapse type are provided in **Supplementary Table 3**.

#### Plasticity Protocols

The plasticity simulations were performed by increasing the conductance values of different types of synapses in the microcircuit model. Specifically, the three conditions tested were PYR plasticity, (PYR-LTP) which involved a 50% increase in the CI-AMPA conductance value of PYR to PYR connections, PVI plasticity (PVI-LTP), which involved a 50% increase in the CP-AMPA conductance value of PYR to PVI connections, and PYR and PVI plasticity (PYR+PV-LTP), which involved a 50% increase in both the CI-AMPA and CP-AMPA conductance values of PYR to PYR and PYR to PVI connections (see also **Supplementary Table 3**).

#### Microcircuit Configuration

The biologically constrained CA3 microcircuit model comprised 22 neurons, specifically 20 PYRs and 2 PVIs, maintaining an excitation to inhibition ratio of 1/10. In each random simulation trial (n=10), each PYR contacted up to 4 other PYRs with 1 CI-AMPA and 1 NMDA synapse activation per contact (convergence=1). On the other hand, every PV BC received synaptic input (convergence=1) from 5 different PYRs in each simulation trial. Additionally, every PYR received 1 feedback inhibitory GABAergic input from each PV BC per simulation trial. Each of the 2 PVIs formed 4 GABAergic synapses per simulation trial and was self-inhibited through autapses. To record the local field potential (LFP), an *in silico* electrode was simulated based on NEURON’s extracellular function (based on^73^). The electrode was placed close to the PYR somata and remained in the same position throughout the simulation trials. The sampling frequency was set at 10 kHz. For simplicity reasons, other inhibitory interneuron types such as dendrite-targeting interneurons and network properties, such as gap junctions, were not simulated as they were not relevant to the experimental observations of this study.

#### Inputs and Simulation

The input was modeled as an artificial presynaptic population using NEURON’s NetStim function (interval=30, number=30). The input targeted only the PYR population of the microcircuit for 1 second, and 3 input synapses were activated in every pyramidal cell (see **Supplementary Table 3**), mimicking the experimental protocol of KA activation primarily in the pyramidal cells. Additionally, both pyramidal cells and PV BCs populations received the same excitatory subthreshold background fluctuation input (subthreshold noise). For every case (including baseline, PYR-, PVI-, or PYR+PVI-LTP), we ran the microcircuit for 10 random trials. In every trial, the number of total synaptic contacts and the connectivity ratios remained identical, but different random neurons were connected to different random neurons. To capture any variability related to morphological features, synapses were allocated in different random dendrites and locations across the selected dendrites in every trial. The recording time was 2 seconds for every simulation trial.

### Data Analysis and Statistics

#### LFP recordings

Recordings were band-pass (1 – 100 Hz, 12 pole IIR digital filter) and band-stop filtered (49 – 51 Hz, 12 pole IIR digital filter). Power spectra were calculated every 30 seconds throughout the recording in each recorded slice individually. Peak power and peak frequency were determined offline by using custom-made MATLAB scripts and visualized as Time-Power- and Time-Frequency-plots, respectively. Slices were excluded from analysis if 1) they only displayed synchronous activity for an interval shorter than ten minutes or during only one induction period or 2) network activity was instable, e.g. peak power would intermittently decrease during KA-application by > 10%.

For each included slice, a 10-minute time window was detected corresponding to maximal peak power in the first induction period. Peak power and frequency were extracted from an additional 10-minute power spectral density obtained with MATLABs pwelch function (0.34 Hz resolution) and an auto-/cross-correlograms obtained over the same interval with the xcorr function. For experiments investigating network activity in two separate episodes of KA-application, the identical analysis was performed twice in time control to the first application period.

#### pMEA recordings

Recordings were pre-processed and peak power and frequency values as well as cross-correlograms for each channel were obtained identically to LFP recordings. Channels included in subsequent analysis were identified by their positioning below the pyramidal cell layer, marked as nodes in relation to their position on the pMEA which were connected via edges to the respectively adjacent selected electrodes (MATLAB digraph function, interelectrode distance 100 µm). For each individual recording, a “lead electrode”, corresponding to the highest peak power value determined, was identified. Intralaminar distances to this electrode were calculated with the “shortestpath” MATLAB function.

#### Simulation data

Data were band-pass and band-stop filtered as LFP recordings. Power spectra were obtained from the entire 2-second recording with the same parameters as LFP recordings and peak power values extracted between 20 – 40 Hz.

#### Statistical analysis

Statistical analysis and data visualization were performed in R 3.5.0. Either Student’s t-test, the Mann-Whitney U-test or Wilcoxon’s Signed Rank test (all two-tailed) were used for statistical comparison of unpaired or paired samples, respectively. The Bonferroni-Holm method was applied to p-values in group comparisons. Kendall’s Signed Rank test was used for correlation analysis. Significance was set to an α-level of 0.05. Data are presented as Mean ± SEM or, in the case of peak power values, Median [1st quartile, 3rd quartile]. Crosses in boxplots denote the mean. Numbers in barplots or below boxplots denote the sample size with n referring to the number of slices tested or simulation trials, respectively.

## Contributions

M.D.H. and J.R.P.G. conceived the study. M.D.H. designed and performed *ex vivo* experiments, data analysis and created the first manuscript draft. A.T. designed and implemented the computational model. All authors contributed to the final manuscript.

## Code and Data availability

Experimental datasets and analysis scripts can be provided by the authors upon reasonable request. The computational model’s source code and the datasets used to generate Figure 4 are publicly available at www.github.com/AlexandraTzilivaki.

## Acknowledgments

Carmen Birchmeier and James Poulet kindly provided PV-Cre and Ai9 animals, respectively. We thank Peer Wulff for the generation and provision of loxp-mG5 mice. Benjamin Rost kindly provided Flex-hM4Di mice. Luna Soso Zradkovic kindly assisted with imaging. Andrea Wilke performed technical assistance with animal husbandry. We thank Benjamin Rost, Nikolaus Maier and Jonas Sauer for insightful comments on an earlier version of the manuscript. MDH and JRPG are supported by the Deutsche Forschungsgemeinschaft (DFG, German Research Foundation, TRR 295 Project ID 424778381 and FOR 2143 Project ID 245861656). DS is supported by the Einstein Foundation Berlin, by the European Research Council (ERC) under the Europeans Union’s Horizon 2020 research and innovation program (BrainPlay Grant, Agreement No. 810580), by the DFG, SFB-958 – Project ID 184695641, Project ID 431572356, FOR 3004 – Project ID 415914819, SFB 1315 – Project ID 327654276 and under Germany’s Excellence Strategy – Exc-2049-390688087 NeuroCure), and by the Federal Ministry of Education and Research (BMBF, SmartAge – Project ID 01GQ1420B). AT is supported by the DFG with the SFB 1315-A01 ‘Brenda Milner Award 2023’, and by a PhD Fellowship from the Einstein Center for Neurosciences Berlin.

---

**Supplementary Figure 1.**
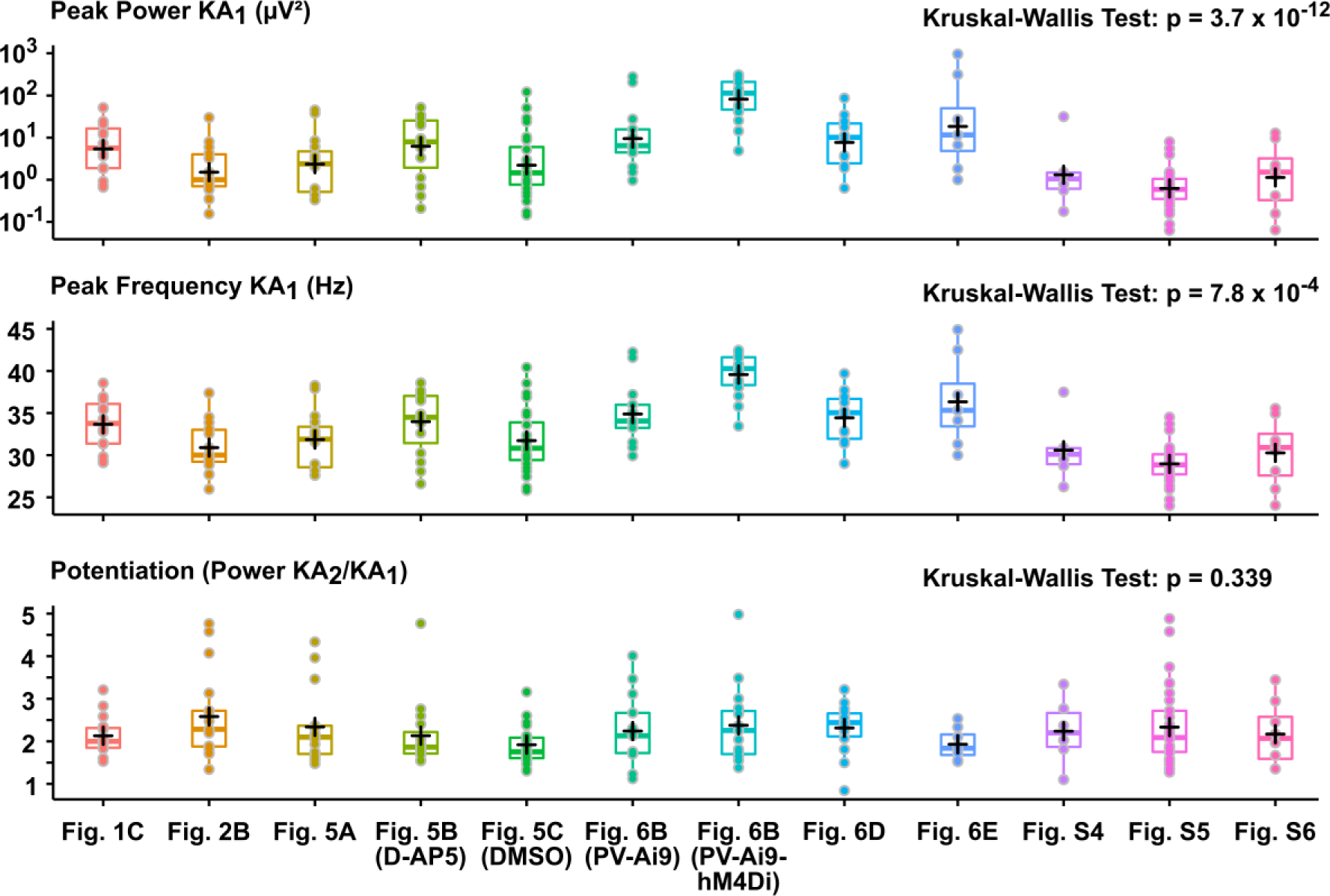
Multi-trial stability of gamma-potentiation in LFP recordings. Boxplots of all control experiments in the twelve interventions reported. Crosses denote the mean. *Above*: Log-scale boxplot of peak gamma-power in the first induction period (KA1). *Centre*: Peak frequency of oscillations in the first induction period. *Below*: Magnitude of gamma-potentiation. Note the high variability of KA1 peak power and frequency amongst trials, which is not mirrored in subsequent potentiation of peak power.

**Supplementary Figure 2.**
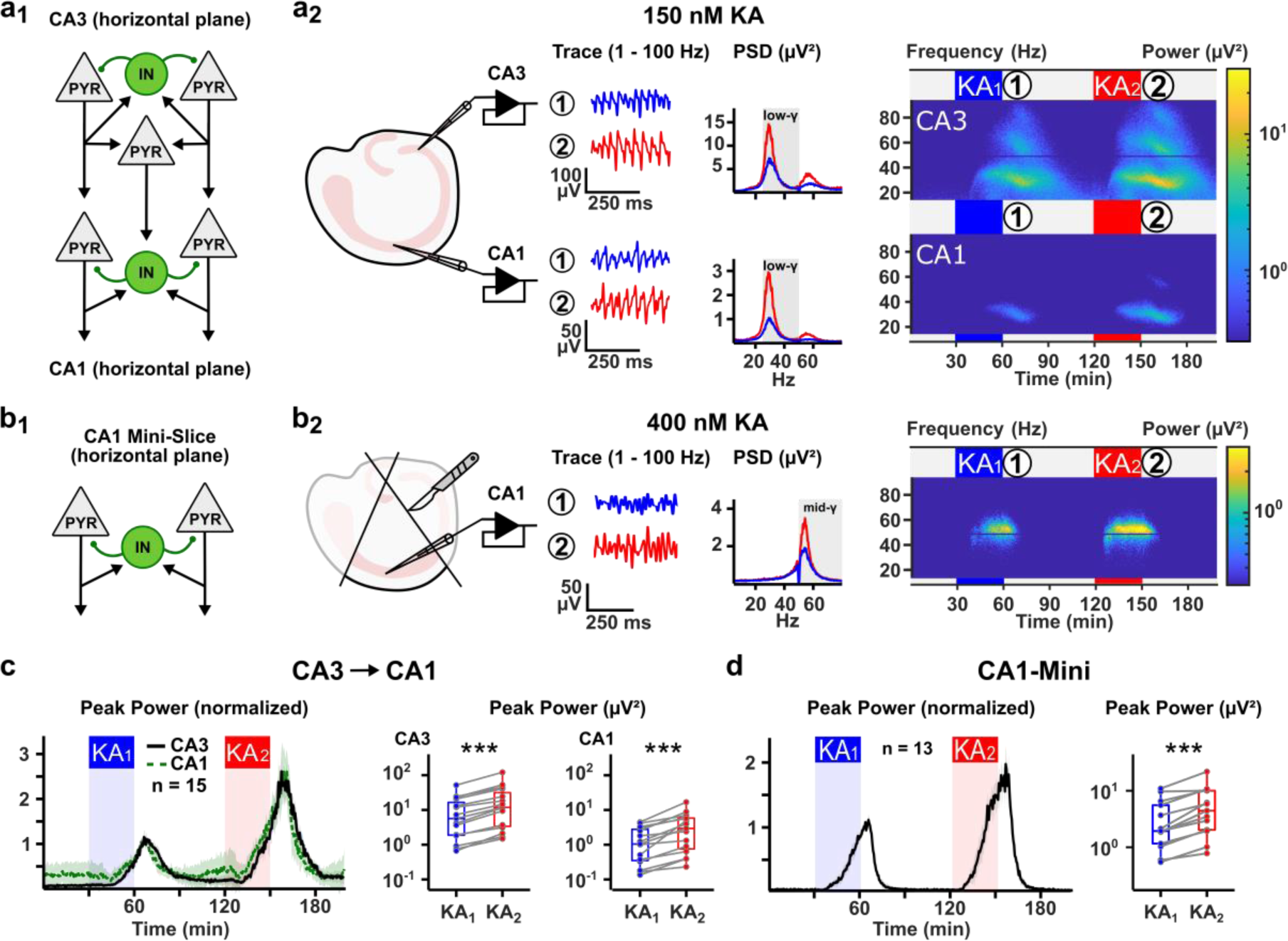
Gamma-potentiation in paired CA3-CA1 and CA1-Mini LFP recordings. **a1** Schematic of CA3 and CA1 connectivity in the horizontal slice plane. CA3 pyramidal cells (PYR) are interconnected amongst themselves, local interneurons (INs) and CA1 PYRs and INs via the Schaffer collateral pathway. CA1 PYRs do not connect amongst each other but are interconnected with local CA1 INs. **a2** *Left:* Schematic of a hippocampal slice with LFP electrodes in CA3 and CA1. *Centre:* Exemplary experiment as in Fig. 1A with simultaneous recordings of CA3 and CA1. *Right:* Pseudocolor plots of the entire experiment. **b1** Schematic of the CA1-Mini microcircuit, lacking input from CA3. **b2** *Left*: Schematic drawing of the hippocampus with surgical cuts separating CA1 from the subiculum and CA3. A recording pipette was placed in stratum pyramidale. *Centre and Right*: same as in **a2** but for an exemplary recording in a CA1-Mini slice. Gamma-activity was induced with 400 nM KA instead of 150 nM. Grey inset “mid-gamma” in the PSDs denotes the window spanning 50 – 80 Hz. **c** *Left* Time-Power plot for all paired CA3-CA1 recordings normalized to the first induction period. *Right* Paired boxplots of peak gamma-power in both application periods in CA3 and CA1. **d** Same as **c**, but for the experiments in CA1-Mini slices. *** denotes p < 0.001 (Wilcoxon Signed-Rank test).

**Supplementary Figure 3.**
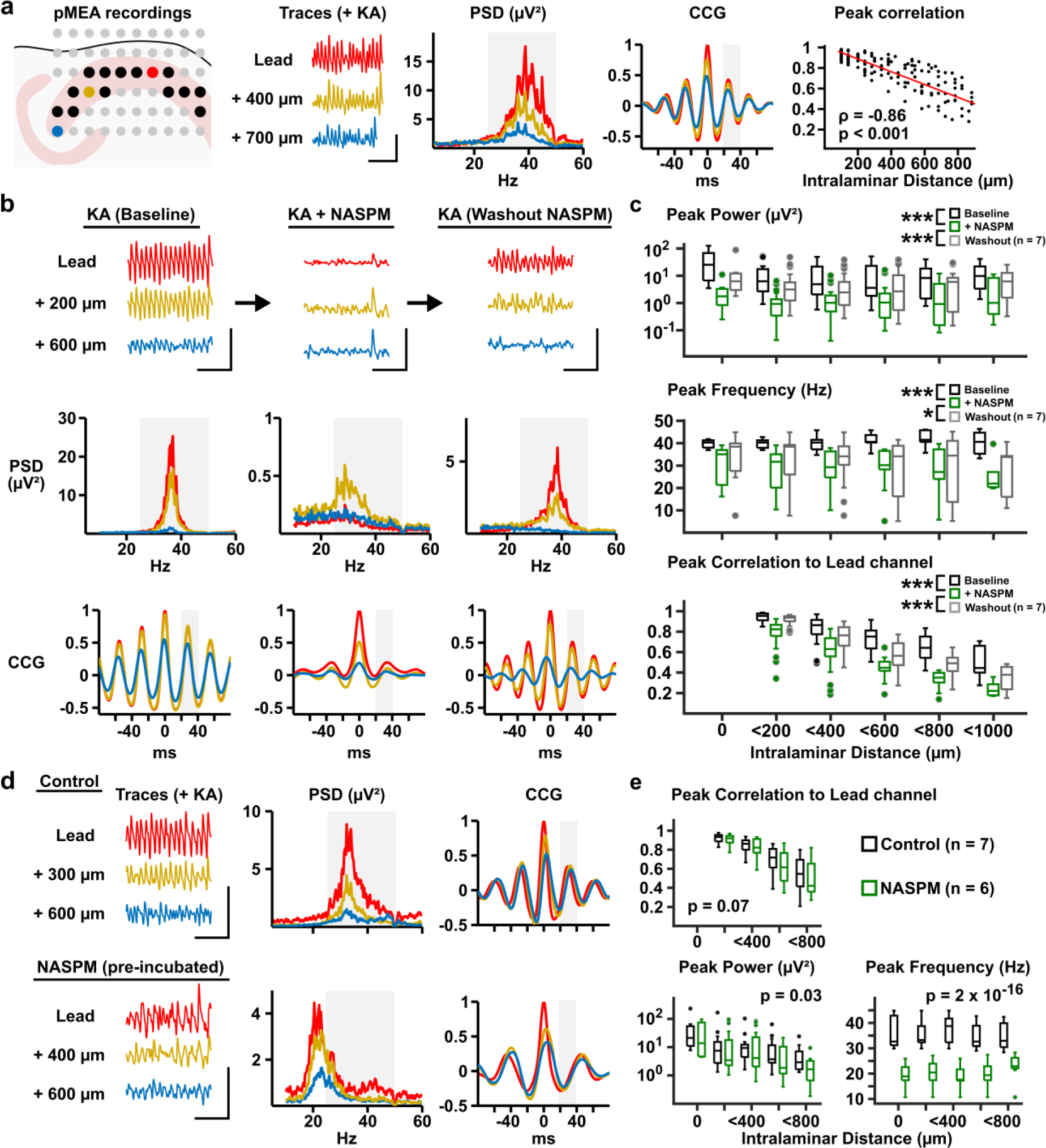
CP-AMPARs generate and maintain the gamma-rhythm in pMEA recordings. **a** Gamma-oscillations are generated across CA3 with decreasing inter-site cross-correlation. *Left, Centre*: Exemplary recording configuration in the 6 × 10 MEA grid with selected electrodes in black. Red, yellow and blue electrodes resemble the recordings displayed in the traces, PSDs and cross-correlograms (CCG) after application of KA (200 nM). The red correlogram represents the auto-correlogram of the lead electrode (highest power in PSD). *Right*: Scatter plot of pooled peak cross-correlation of electrodes to the lead electrode. A linear fit is superimposed. Rho and p refer to the results of Pearson’s correlation test. **b** Gamma-oscillations are maintained by CP-AMPARs. An exemplary experiment with traces at three sites (above) and the corresponding PSDs (centre) and correlograms (below). After an initial induction of gamma-oscillations with 200 nM KA (left), NASPM is added to the bath (centre) and oscillations largely dissipate. Following washout of NASPM (right), oscillations re-appear in 2/3 sites. **c** Boxplots to the experiments in **b**. Values are displayed for peak power, frequency and cross-correlation as a function of intralaminar distance. *** and * denote p < 0.001 and < 0.05, respectively. P-Values obtained from paired T-tests adjusted with the Bonferroni method. Peak power values were log-transformed before statistical testing. **d** CP-AMPAR lacking rhythms are similar to gamma-oscillations in power and inter-site cross-correlation, but markedly reduced in peak frequency. Exemplary experiments as in **a** for a control slice (above) and a slice pre-incubated with NASPM (50 µM) before application of KA (200 nM). In the presence of NASPM, the peak of the PSD and correlograms is clearly below the low gamma-frequency range (25 – 50 Hz and 20 – 40 ms for PSD and correlograms, respectively). **e** Boxplots of recorded peak values for power, frequency and cross-correlation to the lead channel as a function of intralaminar distance toward the lead channel. A clear reduction in peak frequency is observed across all recording sites after pre-incubation with NASPM. P-Values obtained from respective two-way ANOVAs.

**Supplementary Figure 4.**
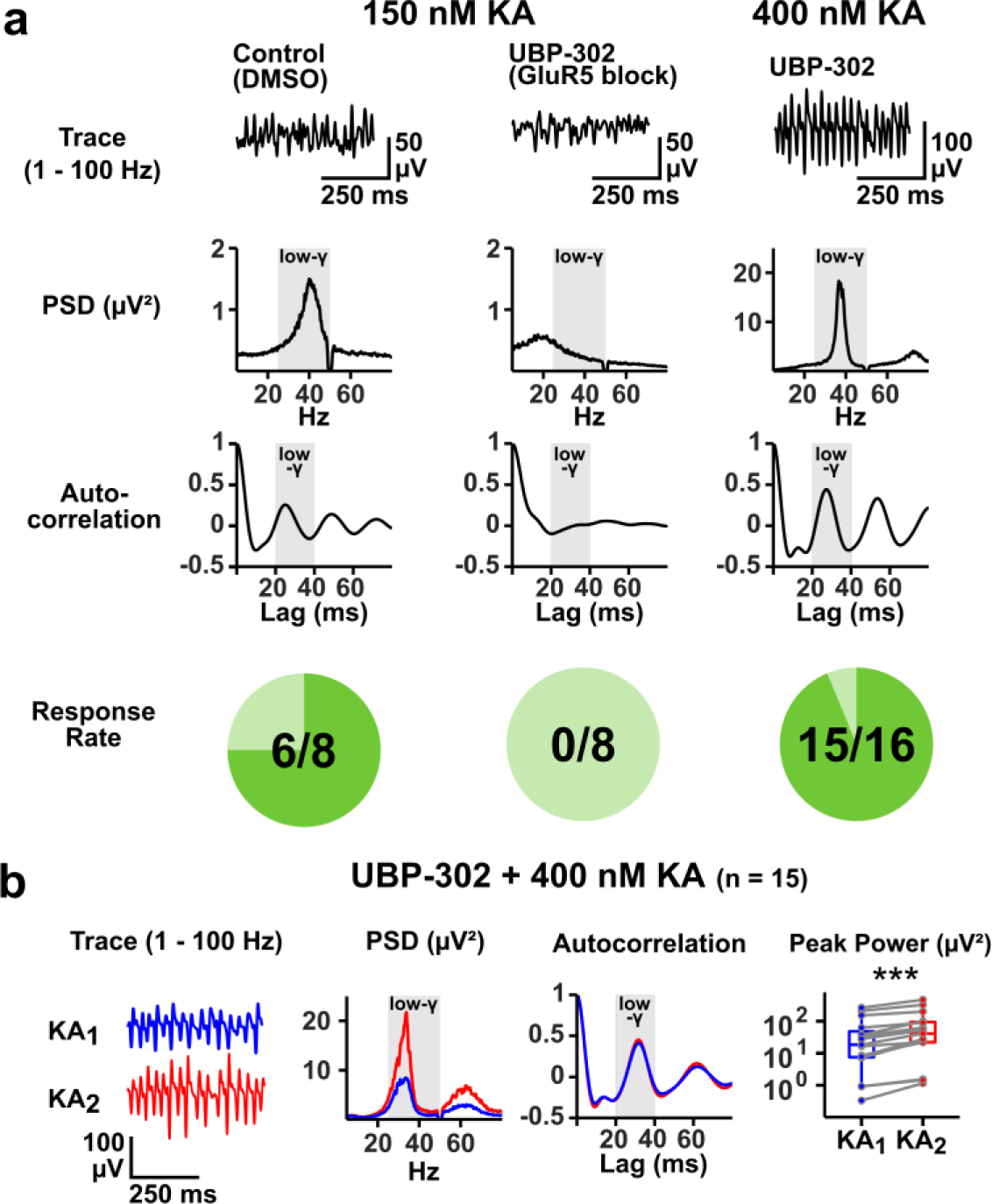
Blockade of GluK1 with UBP-302 attenuates the induction of gamma-oscillations but does not alter gamma-potentiation. **a** Exemplary traces, power spectra and ACGs for DMSO Control experiments and UBP-302 pre-incubation with two concentrations of KA (150 and 400 nM). At 150 nM KA, pre-incubation with UBP-302 prevents the generation of a discernible rhythm in the ACG. *Below*: Pie charts of the response rates of the individual paradigms. **b** Gamma-potentiation in the presence of UBP-302 at 400 nM KA. *Left*: Exemplary traces during the first and second induction period (KA1 and KA2, respectively). *Centre*: Corresponding PSD and ACG. Grey insets “low-gamma” in the PSDs and ACGs denotes the window spanning 25 – 50 Hz. *Right*: Paired boxplot of peak gamma-power during both induction periods. *** and ** denote p < 0.001 and p < 0.01, respectively (Student’s t-test corrected by Bonferroni-Holm in barplots, Wilcoxon Signed-Rank in boxplots).

**Supplementary Figure 5.**
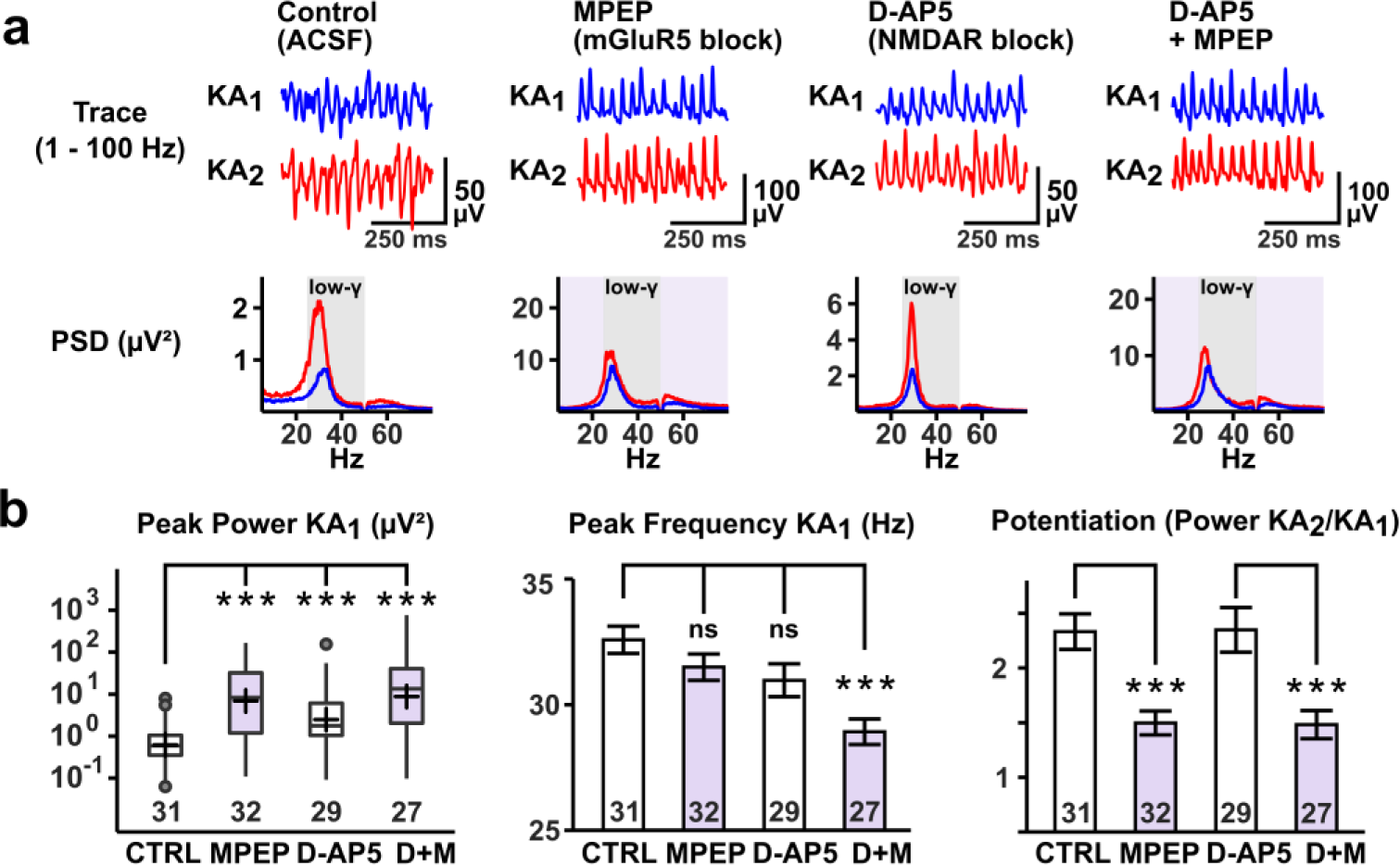
Supplementary Figure 5 mGluR5, not NMDARs, contributes to gamma-potentiation. Blockade of NMDARs and mGluR5 increases peak gamma-power and decreases peak gamma-frequency. **a** Exemplary traces and power spectra for Control, D-AP5, MPEP and D-AP5 and MPEP conditions during both induction periods (KA1, KA2). **b** Summary statistics for all conditions. *Left*: Boxplot of peak gamma-power in the first induction period for all conditions. *Centre*: Barplot of peak gamma-frequency during the first induction period. *Right*: Barplot of potentiation of peak power in all conditions. *** and ns denote p < 0.001 and p > 0.05 (Student’s t-test corrected by Bonferroni-Holm in barplots, Wilcoxon Signed-Rank in boxplots). Purple insets denote the application of MPEP.

**Supplementary Figure 6.**
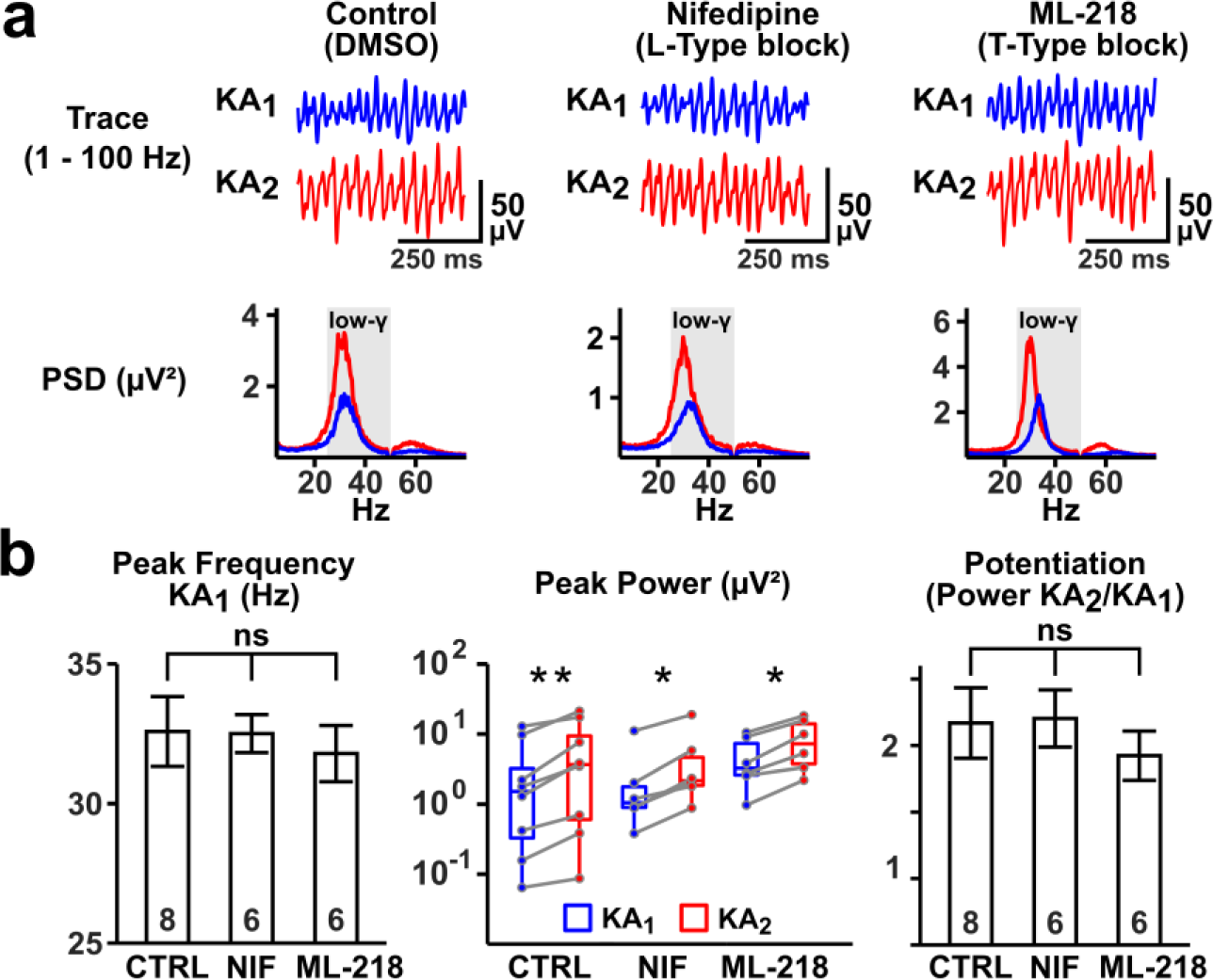
Blockade of L- or T-Type voltage-gated calcium channels does not affect CA3 gamma-activity or subsequent potentiation. **a** Exemplary traces and PSDs for DMSO control, nifedipine and ML-218 conditions during both induction periods (KA1, KA2). **b** Summary statistics for all conditions. *Left*: Barplots of peak gamma-frequency during the first induction period. *Centre*: Paired boxplots of peak gamma-power for both induction periods in all conditions. *Right*: Barplot of potentiation of peak power in all conditions. **, * and ns denote p < 0.01, p < 0.05 and p > 0.05 (Student’s t-test in barplots, Wilcoxon Signed Rank test in boxplots).

**Supplementary Figure 7.**
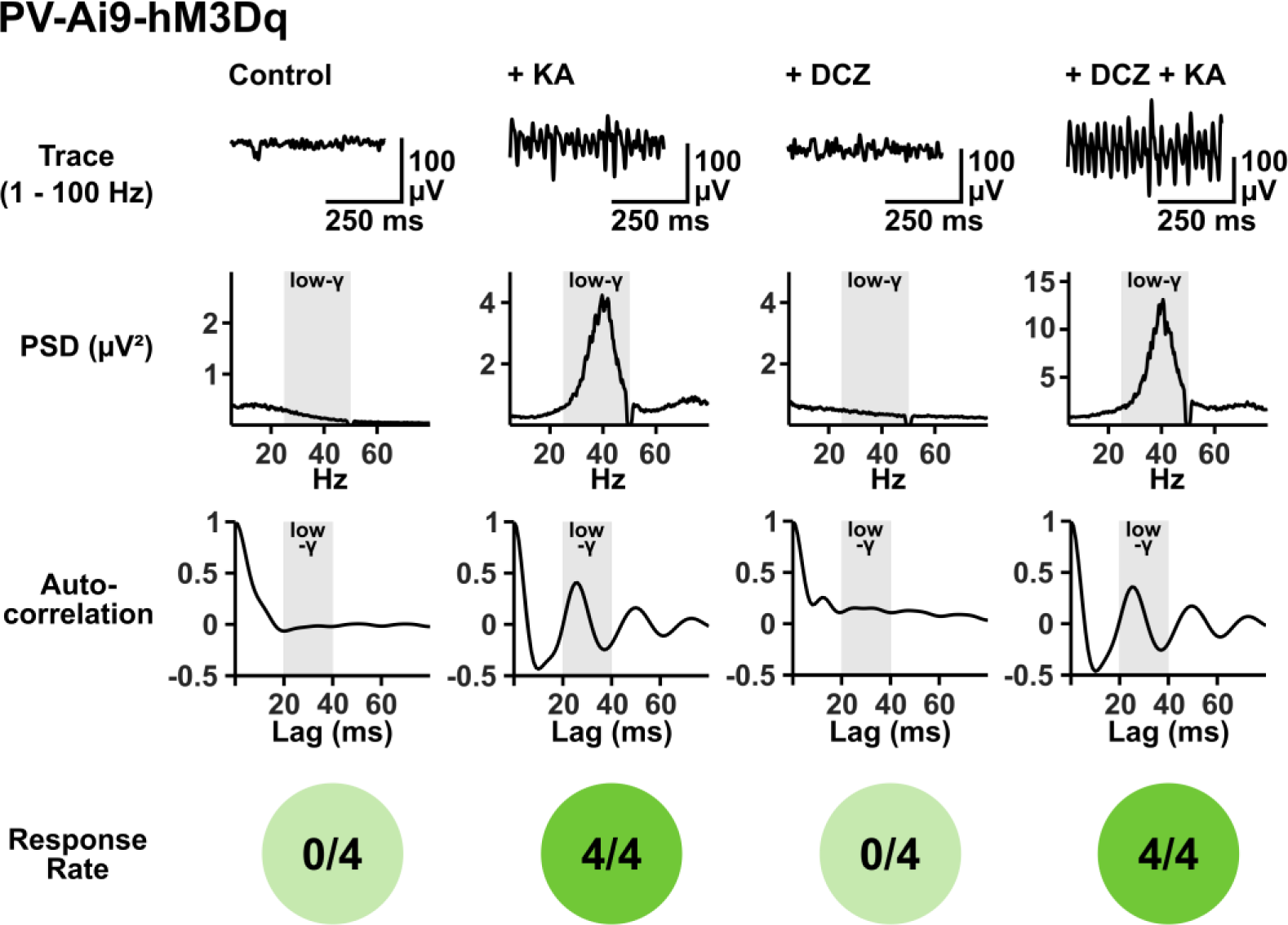
Sole activation of hM3Dq in PVIs is insufficient to synchronize CA3 network activity. Control experiments for sole PV-hM3Dq activation. Traces, PSDs and ACGs of conditions testing whether gamma-activity can be induced with just 3 µM DCZ. Pie plots below indicate the numbers of slices in which oscillations were induced for each condition. KA (150 nM) or DCZ were applied or co-applied for 30 minutes.

**Supplementary Table 1.** Passive parameters and active conductances values for all compartments of the pyramidal model cell.

| Mechanism | Soma | Proximal<br>Apical<br>Dendrite | Distal<br>Apical<br>Dendrites | Basal<br>Dendrites |
| --- | --- | --- | --- | --- |
| Leak conductance [S/cm<sup>2</sup>] | 0,0002 | 0,0002 | 0,0002 | 0,0002 |
| Na<sup>+</sup> conductance [S/cm<sup>2</sup>] | 0,0105 | 0,0084 | 0,0084 | 0,0084 |
| Delayed rectifier K<sup>+</sup> conductance [S/cm<sup>2</sup>] | 0,00086 | 0,00086 | 0,00086 | 0,00086 |
| Proximal A-type K<sup>+</sup> conductance [S/cm<sup>2</sup>] | 0,0075 | 0,015 | - | 0,0075 |
| Distal A-type K<sup>+</sup> conductance [S/cm<sup>2</sup>] | - | - | 0,04875 | - |
| M-type K<sup>+</sup> conductance [S/cm<sup>2</sup>] | 0.06 | 0.06 | - | 0.06 |
| I<sub>h</sub> conductance [S/cm<sup>2</sup>] | 0.00005 | 0.0001 | - | 0.00005 |
| L-type Ca<sup>2+</sup> conductance [S/cm<sup>2</sup>] | 0.0007 | 0.00003 | - | 0.00003 |
| R-type Ca<sup>2+</sup> conductance [S/cm<sup>2</sup>] | 0.0003 | 0.00003 | - | - |
| T-type Ca<sup>2+</sup> conductance [S/cm<sup>2</sup>] | 0.00005 | 0.0001 | - | 0.0001 |
| Ca<sup>2+</sup>-dependent sAHP K<sup>+</sup> conductance [S/cm<sup>2</sup>] | 0.0015 | 0.001 | - | 0.0005 |
| Ca<sup>2+</sup>-dependent mAHP K<sup>+</sup> conductance [S/cm<sup>2</sup>] | 0,9 | 0,03 | - | 0,08 |
| Membrane capacitance C<sub>m</sub> [μF/cm<sup>2</sup>] | 1 | 1 | 1 | 1 |
| Membrane resistance R<sub>m</sub> [Ohm cm<sup>2</sup>] | 6000 | 6000 | 6000 | 6000 |
| Axial resistance R<sub>a</sub> [Ohm cm] | 150 | 150 | 150 | 150 |

**Supplementary Table 2.** Electrophysiological characterization of the pyramidal model cell.

| Parameter | Value |
| --- | --- |
| Rheobase (pA) | 200 |
| Input Resistance (MΩ) | 142 |
| Spike Threshold (mV) | -43,11 |
| Spike Amplitude (mV) | 73,11 |
| Spike Overshoot (mV) | 30 |
| Resting Membrane Potential (mV) | -70 |
| AP Frequency (Hz) during Rheobase current clamp | 6 |

**Supplementary Table 3.** CA3 microcircuit synaptic conductance values.

| Synapse Type | Pyramidal model | PV Basket Cell model |
| --- | --- | --- |
| INPUT AMPA | 0,002 | - |
| INPUT NMDA | 0,005 | - |
| CI-AMPA | 0,0017 (baseline) | - |
|  | 0,00255 (plasticity) |  |
| CP-AMPA | - | 0,0015 (baseline) |
|  |  | 0,00225 (plasticity) |
| NMDA | 0,0051 | 0,0016 |
| GABA | 0,004 | 0,005 |
| Autapse GABA | - | 0,007 |

